# A geothermal amoeba sets a new upper temperature limit for eukaryotes

**DOI:** 10.1101/2025.11.24.690213

**Authors:** H. Beryl Rappaport, Natalie A. Petek-Seoane, Tomáš Tyml, Felix Mikus, Kurt LaButti, Godwin Ani, Jessica K. Niblo, Ethan MacVicar, Rachel M. Shepherd, Ignacio de la Higuera, Samuel J. Lord, Gautam Dey, Gordon V. Wolfe, Omaya Dudin, Shahar Sukenik, Laura A. Katz, Kenneth M. Stedman, Kristen Skruber, Frederik Schulz, R. Dyche Mullins, Angela M. Oliverio

## Abstract

The study of temperature limits has transformed our knowledge of the boundaries of life but has been largely focused on bacteria and archaea. We isolated a novel geothermal amoeba, *Incendiamoeba cascadensis*, that divides at 63°C (145.4°F), establishing a new record for the upper temperature limit across all eukaryotes. We demonstrated cellular proliferation with growth experiments and visualized mitosis *via* expansion microscopy. Using high-temperature live-cell imaging, we quantified movement up to 64°C. We assembled the genome of *I. cascadensis* and using comparative genomics found an enrichment of genes related to proteostasis, genome stability, and sensing the external environment. Taken together, our findings challenge the current paradigm of temperature constraints on eukaryotic cells and reshape our understanding of where and how eukaryotic life can persist.

## Main Text

The thermal limits of life on Earth have long been a source of scientific fascination and discovery: by studying the physical factors at the limits of life, we can broaden our understanding of constraints on cellular biology and of where life can emerge and persist. Temperature regulates life from enzyme stability, to cellular functions and metabolic processes, to the distribution of species across ecosystems (*1*, *2*) and impacts physical parameters such as pH, viscosity, density, and solubility (*1*). Characterization of bacteria and archaea that inhabit high temperature environments such as geothermal springs and hydrothermal vents has uncovered some of Earth’s most metabolically and phylogenetically diverse organisms, including most of the candidate archaeal phyla (*2*, *3*).

Thermophiles are organisms with high thermal limits and optimal growth at or above 45°C (*4*). Thermal limits are defined by three thresholds: completion of the life cycle (e.g. cellular replication), metabolic activity (e.g. motility and feeding), and survival (*1*). The current highest known temperature threshold is 122°C, at which only the archaean *Methanopyrus kandleri* can grow (*5*). Among bacteria, *Geothermobacterium ferrireducens* can grow at temperatures up to 100°C (*6*). These growth temperatures are remarkable given the challenges associated with thermal stress, including increased likelihood of protein and other biomolecule denaturation and impacts on membrane fluidity (*2*, *7*).

Surprisingly, parallel efforts to characterize eukaryotic thermophiles and define upper temperature thresholds remain extremely limited (*8–11*). The current upper temperature limit of growth for any eukaryotic organism is 60°C, at which a few species of fungi and red algae can replicate (*12–16*). Other eukaryotes have been documented growing up to 57°C, including the amoeba *Echinamoeba thermarum* (*17*), but the vast majority of eukaryotic diversity in high temperature environments remains unexplored (*8*). Thermophilic red algae and fungi offer some insight on high temperature adaptations including coordinated signaling pathways for efficient proteostasis and chaperone production (*18*, *19*). However, both the molecular mechanisms that could inherently limit eukaryotic growth at a specific temperature and the true upper temperature limit for eukaryotic cells remain unknown. Although 62°C has been hypothesized as a theoretical upper bound based on a presumed lack of organelle membrane thermostability, this was from a limited number of samples that did not yield fungal growth above 60°C (*12*).

Here, we report the discovery of a geothermal amoeba isolated from Lassen Volcanic National Park (LVNP; California, USA) and (i) describe its evolutionary placement, morphology, and biogeography; (ii) determine its maximum temperature for replication and motility; and (iii) shed light on the cellular and molecular strategies used to protect cellular structure and function in extreme heat. Our findings establish new upper temperature limits for cellular replication and motility for a eukaryotic cell.

### A novel genus of amoeba cultivated from geothermal springs

We isolated a novel amoeba from 14 out of 20 geothermal sampling locations over a three year period (2023–2025) along a tributary of Hot Springs Creek in LVNP. Tributary temperatures ranged from 49°C-65°C and pH from 6-7 (**Fig. S1A-C; Table S1**). Site temperatures and geochemistry are highly stable over decades (*20*). Samples yielding the novel amoeba were from water, biofilm, and sediment. To obtain cultures, we enriched samples with a wheatberry to stimulate growth of the *in situ* bacterial community, which in turn enriched the growth of the bacterivorous amoeba (**Fig. S1D**). Within one to two weeks post enrichment, amoebae were visible from incubations at 57°C and 60°C, and within three weeks, amoebae were visible at 63°C. We also isolated a strain of *Vermamoeba vermiformis* at room temperature (**Table S1; Fig. S1B**) and used it for morphological comparison. This tributary is one of the few non-acidic geothermal features in LVNP, a vapor-dominated acid-sulfate thermal system (*20*, *21*). Microbial communities at Boiling Springs Lake, an acid-sulfate geothermal feature about one kilometer from this study site, have been characterized (*21*, *22*), but this site was not included in previous efforts.

To determine the phylogenetic placement of this novel amoeba, we obtained two replicate near-complete genomes from cultures (**Fig. S1B**) and used the phylogenomics pipeline EukPhylo (*23*). Briefly, we produced a phylogeny from a multi-sequence alignment of 102 concatenated genes. The amoeba falls within Tubulinea (Amoebozoa) (**Fig. 1A**) in the clade Echinamoebida, which notably contains *Echinamoeba*, the genus that includes the current highest reported amoeba growth temperature. The novel amoeba is sister to *V. vermiformis*, a thermotolerant amoeba often sampled from hot water heaters, capable of growth at room temperature, and with an upper growth limit near 40-45°C (*24*, *25*), 20°C colder than the novel isolate. We built an 18S rRNA tree which supports this placement (**Fig. S2**). The 18S rRNA sequence of the novel amoeba is 94% identical to the room-temperature species *V. vermiformis*, in contrast to less than 1% 18S sequence divergence within the *Vermamoeba* genus (*26*), and the branch length distance between the novel amoeba and *V. vermiformis* was outside a 95% confidence interval from six other within-genus pairs of species (**Table S2**). Coupled with the morphological and ecological evidence described below, we propose the novel amoeba as a new genus *Incendiamoeba* (Latin incendarius meaning igniting fire, “fire amoeba”) and species *cascadensis*; coming from the Cascades, the mountain range where the amoeba was discovered.

**Fig. 1.**
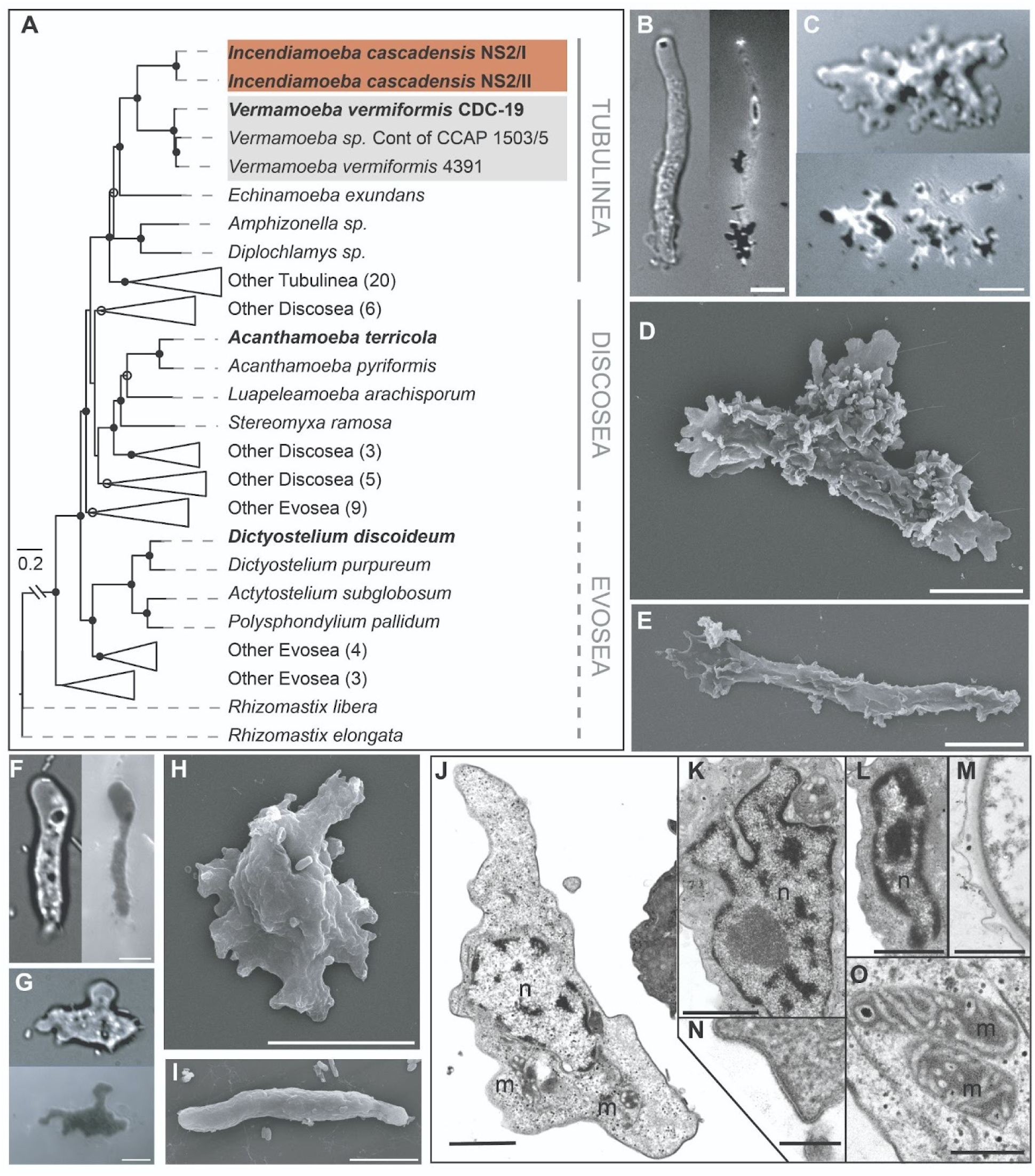
Phylogenetic, morphological, and physiological data support establishment of *Incendiamoeba cascadensis*, a novel genus and species within Amoebozoa. (**A**) Concatenated tree of 102 Amoebozoa genes, nodes labeled with filled circles have 100% bootstrap support, open circles have >90% bootstrap support. Novel amoeba sequences are highlighted in dark orange, *Vermamoeba vermiformis* in grey, and the species used for genome comparison are bolded. (**B**) DIC and corresponding RICM of representative *I. cascadensis* in vermiform state and (**C**) in amoebiform state. (**D**) SEM of *I. cascadensis* in amoebiform state and (**E**) SEM in vermiform state. (**F**) DIC and corresponding RICM of representative *V. vermiformis* in vermiform state and (**G**) in amoebiform state. (**H**) SEM of *V. vermiformis* in amoebiform state and (**I**) SEM in vermiform state. All scale bars **A-H** are 5µm. (**J-O**) TEM ultrastructure of *I. cascadensis* including: (**J**) Overview of fine structure of trophozoite, nucleus (n), mitochondria (m) (**K**) Lobate nucleus with dispersed chromatin aggregates. (**L**) Nucleus with chromatin aggregates beneath the nuclear envelope. (**M**) Bilayered cyst wall. (**N**) Cell surface of trophozoite. (**O**) Mitochondria with non-branching tubular cristae. Scale bars **J-M** are 1µm, **N** is 200nm, and **O** is 500nm.

To find if *Incendiamoeba* has been detected from previous environmental sequencing efforts and to explore its biogeography, we searched 31,093 publicly available metagenomes from diverse environments globally. We found an identical 18S rRNA gene sequence (**Fig. S2**) in a geothermal microbial mat in the Taupō Volcanic Zone (Waikite Valley, NZ), with temperatures ranging from 62°C near the source to 37°C downstream and pH 8 (*27*). Three additional metagenomes from Yellowstone National Park (USA; reported temperatures of 42–90°C) contained sequences in the same clade, two forming a distinct subclade that may represent additional species-level diversity (**Fig. S2**). Notably, *Incendiamoeba* sequences were only detected in geothermal samples, suggesting its distribution is restricted to high-temperature geothermal areas globally.

We next characterized the morphology of *I. cascadensis* (**Fig. 1B-E**) compared to *V. vermiformis* (**Fig. 1F-I**) using high-temperature light microscopy and scanning electron microscopy and of *I. cascadensis* with transmission electron microscopy (**Fig. 1J-O**). We identified two distinct morphological forms: an elongated, vermiform (worm-like) state similar to that previously described for *V. vermiformis*, (**Fig. 1B,E**) and a more compact, amoebiform state, characterized by pseudopodia (**Fig. 1C,D**). *I. cascadensis* are 38.7 ± 9.4μm long in the vermiform state and 23.5 ± 7.9µm long in the amoebiform (**Table S3; Taxonomic Appendix**). Although *V. vermiformis* and *I. cascadensis* cells exhibit comparable motility and switch between similar vermiform and amoebiform morphologies (**Movie S1-S3**), reflection interference contrast microscopy (RICM) reveals differences in the extent of their substrate adhesion. *I. cascadensis* form much smaller and more transient surface attachments than *V. vermiformis* (**Fig. 1B-C** compared to **Fig. 1F-G; Movie S1-S3**). Thus, while their motility and morphology appear similar, the molecular and biophysical details may differ.

*I. cascadensis* are typically uninucleate and, in rapidly moving vermiform cells, the nucleus appears to be actively driven in the direction of travel (**Movie S4**). We observed *I. cascadensis* preferentially feeding on filamentous bacteria (**Movie S5**), employing a protrusive, phagocytic structure that envelops the prey and exerts forces sufficient to bend and fold the bacterial cells, forming tight loops inside the amoeba (**Fig. S3; Supplementary Text**). Metagenomic sequence abundance and morphology indicate this bacterial prey is *Meiothermus ruber*, a filamentous Deinococcota with an optimum growth temperature of 60°C (*28*), aligning closely with the growth optimum we observe for the novel amoeba. At temperatures above 64°C. *I. cascadensis* form compact but irregularly shaped cysts, ranging in diameter from ∼5 to ∼15µm (**Fig. S4**); encystment is a common strategy for both thermotolerant and mesophilic amoebae to become dormant when conditions are unfavorable (*24*). This genus is also notably distinguished from *Vermamoeba* by its ability to thrive at much higher temperatures, as described below.

### Novel amoeba sets a record upper temperature limit in a eukaryotic cell

Remarkably, *I. cascadensis* grows robustly at the previously defined limit for eukaryotes of 60°C (*12–16*) and continues replicating at a maximum temperature of 63°C (**Fig. 2A-B**). To determine the upper and lower temperature limits for cellular replication of *I. cascadensis*, we conducted growth experiments at 17 temperatures from 30°C to 64°C, with four replicate flasks per temperature, observing culture density over seven days (**Table S4**). We validated incubation temperatures with multiple temperature probes (**Table S4**). To set cutoffs for significant growth at each temperature, we estimated slopes of linear models fit to the log of each growth curve. The minimum growth temperature is 42°C, with no growth at 40°C or lower, and optimal growth is at 55-57°C (**Fig. 2A**; **Table S5**), meaning these amoebae are obligate thermophiles.

**Fig. 2.**
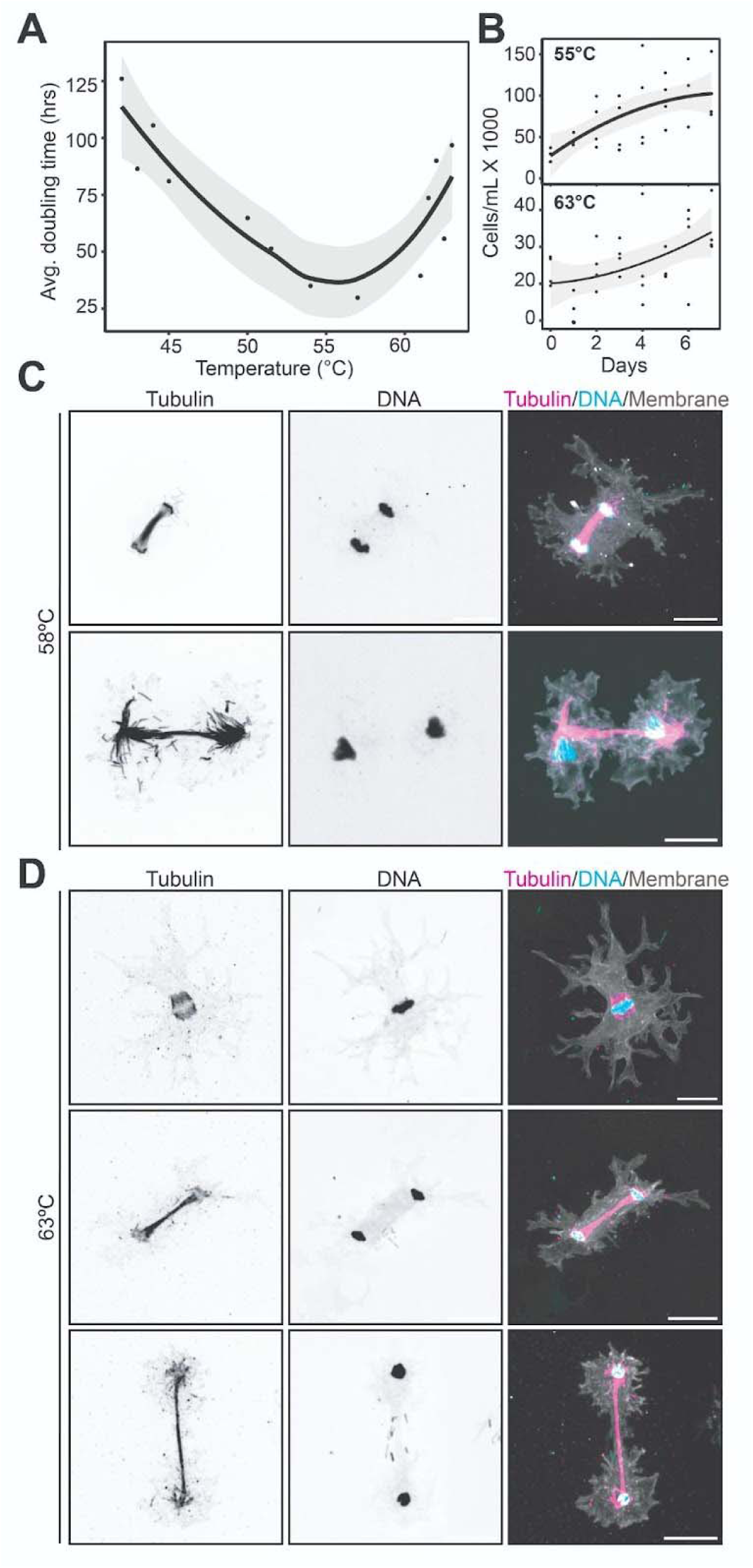
*Incendiamoeba cascadensis* undergoes cellular replication at temperatures up to 63°C. (**A**) Doubling time across temperature range of novel amoeba. Individual points represent average doubling time from the exponential growth period of a seven-day growth experiment. Gray shading represents a 95% CI. (**B**) Example of individual growth curves from 55°C and 63°C. (**C-D**) Ultrastructure expansion microscopy confocal images of mitotic cells fixed at 58°C (**C**) or 63°C (**D**), with tubulin channel (left), DNA (middle), composite (right) with tubulin (pink), DNA (blue), and Bodipy membrane (gray). Top row in **D** consistent with metaphase and in **C** and middle row of **D**, the mitotic spindle array is visible, and cells are arriving at the end of division. The bottom row in **D** displays late telophase. Scale bars 20μm, not adjusted for the expansion factor (approximately 4x expansion).

We also visualized cell division stages (including mitosis) in both cells grown near their optimal range of 58°C and incubated for seven days at their maximum replication temperature of 63°C using ultrastructure Expansion Microscopy (U-ExM), following the protocol described in (*29*). With tubulin (anti-alpha and beta tubulin) and DNA (Hoechst) staining, we identified cells undergoing mitosis at the point of fixation (**Fig. 2C-F**). Metaphase mitotic spindles were barrel-shaped, similar to those observed in the amoebaflagellate *Naegleria gruberi* (*30*), with unfocused poles and no evidence of astral microtubules. In later stages of mitosis, the spindles resemble those of *Dictyostelium discoideum* (*31*). Late anaphase spindles, for example, exhibit both astral microtubules and very long interpolar microtubules in the spindle midzone (*32*).

The temperature dependence of *I. cascadensis* motility exhibits a broad peak, centered at 57°C but extending as high as 64°C, above the previously defined thermal limit for eukaryotes. Briefly, we employed high-temperature, live-cell microscopy (*33*) and automated cell centroid tracking (**Fig. S5**) to characterize motility across a 45°C range (from 25°C-70°C; **Movies S7-S9**). Consistent with our growth curve measurements, we found that cell velocity peaked within the optimal growth range at 57°C (14.5 µm/min; **Fig. 3A**; **Table S6**). To discriminate active motion from passive flow (e.g. advective currents in the media), we included conditions in which all of the cells were encysted (i.e. 25° and 70°C). Due to the random nature of cell crawling, we quantified active motility by plotting mean squared displacement (MSD) versus time (**Fig. 3B; Fig. S6; Supplementary Text**). This method is useful for characterizing trajectories that contain elements of both random and directed motion. In addition to comparing the magnitude of the MSD values at a fixed time point, we also calculated the slopes (α) of log-log plots of MSD versus time. Higher values of α indicate more unconstrained and/or directed motion, while low values indicate frequent periods of immobility. Similar to the velocity and MSD, the value of α peaked between 55-64°C (α > 1, p < 0.0001; **Fig. 3B; Table S7**). In contrast, values of α measured at encystment temperatures were close to zero. Displacement ratios (a measure of trajectory straightness or persistence of motility, **Table S7**) also display a peak in persistence of motion in the 55-64°C range.

**Fig. 3.**
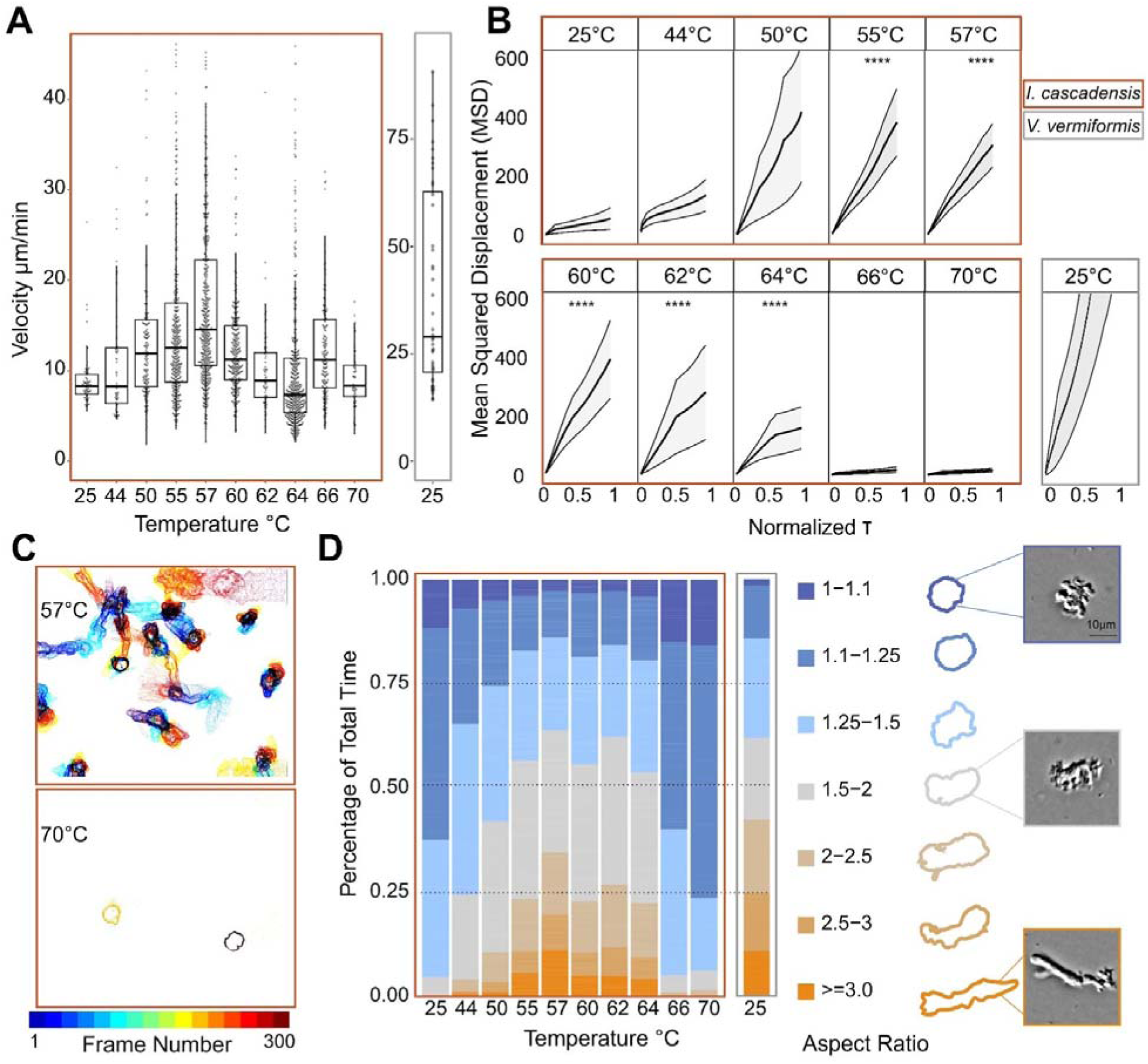
Temperature-controlled live cell imaging reveals multiple motility modes of *Incendiamoeba* up to 64°C. (**A**) Box and whisker plots indicate the median velocity (µm/min) of tracked cell centroids where the percentiles are visualized as follows: 75th (top edge), 25th (bottom edge), and median (bold line); whiskers extend to 1.5×IQR from the box, excluding outliers. Points represent individual particle tracks. (**B**) Mean squared displacement over time of tracked cells across temperature. Error bars represent standard deviation. Asterisks represent significance values (**** = p < 0.0001) for comparisons of α against a reference value of 1 using two one-sided t-tests. Values greater than 1 indicate directed motion. (**C**) Representative contour traces of cells at one second intervals at 57°C and 70°C. Frame number is indicated by color. (**D**) Percentage of total time cells spent at given aspect ratios (>2.5 for vermiform, 1.2-2.5 for amoebiform, 1.0-1.2 for predominantly cysts). Bars represent each temperature recorded and exemplar cell outlines for each aspect ratio are provided along with corresponding DIC images at low, middle, and high aspect ratios. Boxes around **A**-**D** plots are colored by species (*I. cascadensis* in dark orange and *V. vermiformis* in gray).

Similar to other thermotolerant amoebae (*34*, *35*), *I. cascadensis* can recover from encystment at temperatures above its cellular replication limit. At 66°C, cells round up but remain partially active, likely beginning to encyst (**Movies S10-S11**; **Fig. S4, Table S7**). When held at 70°C for five minutes, cells encyst. While they can recover when brought back to 60°C, they are unable to recover from treatment at 80°C. In contrast, *V. vermiformis* cysts are inactivated at 60°C, further differentiating them from *I. cascadensis* (*24*). As *I. cascadensis* live in a stream with fluctuating temperatures, survival across a broad range of temperatures is ecologically relevant.

Motility characterization can also uncover ecological strategies of amoebae, such as space exploration and finding prey. To quantify how shape relates to movement in *I. cascadensis*, we identified cell edges (**Fig. 3C**) and extracted aspect ratios (the ratio of major to minor cell axes; **Table S3**, **Fig. 3D**). Encysted cells exhibited the lowest (i.e. most circular) aspect ratios, while cells at 55-64°C had the highest (i.e. most elongated), peaking at 57°C (**Fig. 3D**). We categorized cells into vermiform, amoebiform, or cyst forms. At 55-64°C, cells spend approximately 30% of their time vermiform, displaying higher velocities (at 57°C: 28.3 µm/min versus 17.1 µm/min; **Table S6**) and straighter trajectories (**Fig. 3D**, **Table S7**). In contrast, amoebiform cells displayed greater trajectory areas, indicating space exploration (57°C mean convex hull: 58.0µm^2^ versus 25.0µm^2^, respectively, **Table S7**).

We observed cells switching frequently between vermiform and amoebiform shapes (**Movie S1, S12**), suggesting that *I. cascadensis* use a two-state motility. On average, *I. cascadensis* switched shapes once every 91.2 ± 3.2 seconds at 57°C while cells at 50°C switched every 235.8 ± 2.1 seconds (**Table S8**). Although this switching has been observed in other amoebae, such as *D. discoideum*, it is much less frequent than *I. cascadensis* (*36*). *I. cascadensis* switched shape 2.6 times more frequently than *V. vermiformis* at their optimal temperatures (**Movie S1, S3, S12, Table S8**). Overall, at 57°C, *I. cascadensis* exhibited lower velocity, explored more space, and changed shape more often than *V. vermiformis* at 25°C. This may be a strategy to alternate foraging and traveling quickly to optimal temperature conditions when living in extreme temperatures that can rapidly fluctuate within a few centimeters (*37*).

### Comparative analyses shed light on high temperature adaptation

To better understand how this amoeba survives at high temperatures, we assembled two high-quality genomes using long-read sequencing and complementary RNA sequencing. Briefly, we assembled genomes of cultures sequenced with PacBio ultra-low input libraries (**Fig. 4A**). To improve gene calls and confirm transcription, we sequenced transcriptomes of two replicate cultures. The genome assembly is 48Mbp and GC content is 36%, slightly larger (contrary to genome reduction in thermophilic fungi (*38*)) and less GC rich than *V. vermiformis* (**Table S9**). Codon usage is distinct from *V. vermiformis* as well (**Fig. 4B**). We recovered a 53kbp mitochondrial genome with 65 predicted genes. The assembly quality metrics were similar to reference genomes included for comparison: *Acanthamoeba terricola* (previously *castellanii*), *D. discoideum*, and *V. vermiformis* (**Table S9**). We estimate assemblies are 83-94% complete based on Benchmarking Universal Single-Copy Ortholog (BUSCO) markers and given shared missing markers among Amoebozoa (*39*)). Beyond offering insight into attributes that may facilitate success, the genomes of *I. cascadensis* also help address the scarcity of protist genomic data (*40*).

**Fig. 4.**
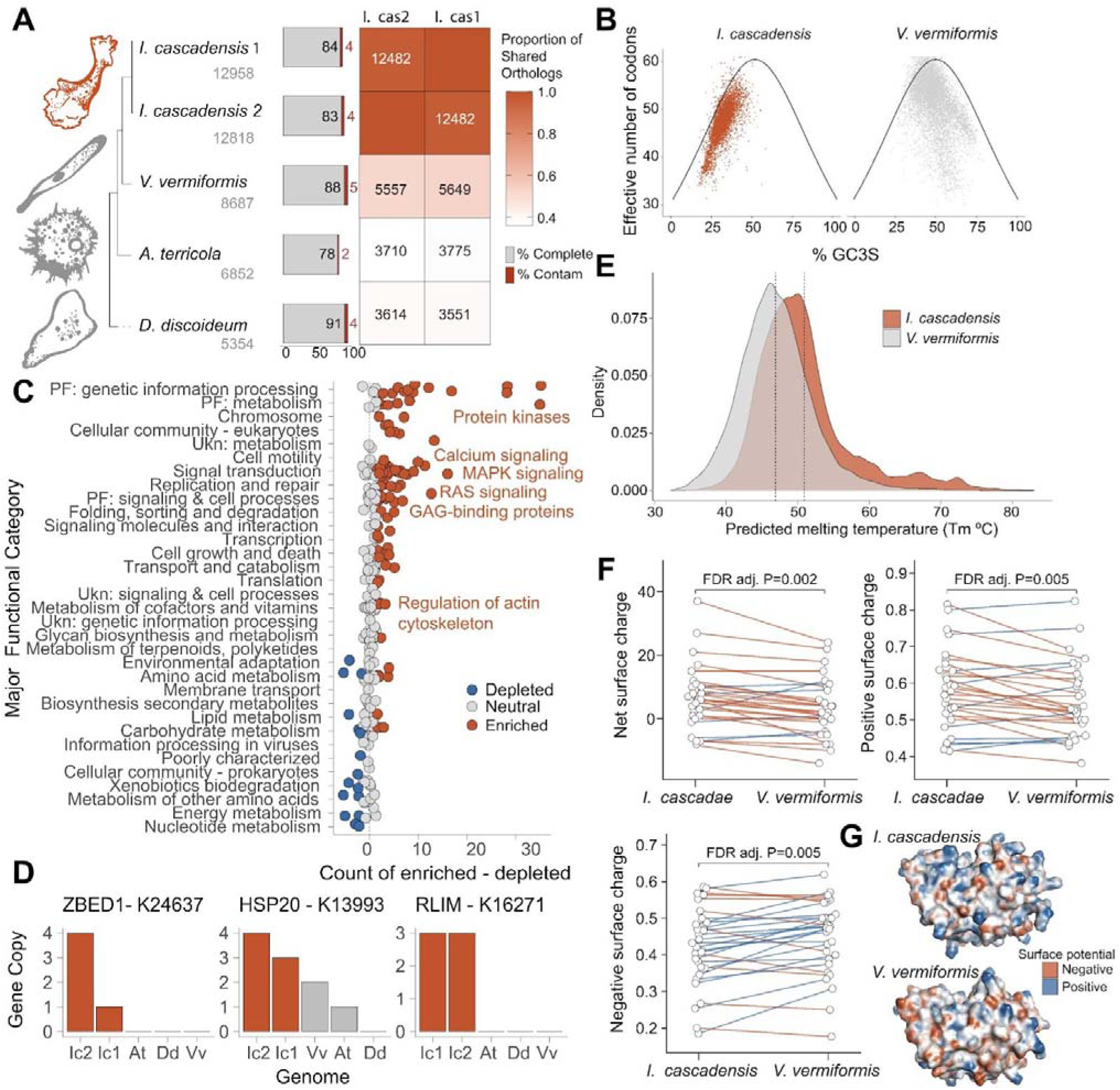
Comparative analyses indicate enrichment of signaling pathways, increased protein stability, and distinct biophysical properties may support thermophilic homeostasis. **(A)** Relationship between two draft *I. cascadensis* genomes and three mesophilic amoebae for comparison (*V. vermiformis*, *A. terricola*, and *D. discoideum*). Total predicted orthologs are below species name. Bars represent percent completeness and contamination. Heat map represents overlap of orthologs between *I. cascadensis* and other amoebae, with darker shading indicating more overlap. (**B**) plots of GC3S (GC content at silent third codon positions) versus effective codon number for *I. cascadensis* and *V. vermiformis*. (**C**) KEGG functions enriched in *I. cascadensis* relative to mesophilic amoebae from **A**. (**D**) Bar plots represent copies of three example genes enriched in *I. cascadensis* (Ic) relative to *V. vermiformis* (Vv), *A. terricola* (At), and *D. discoideum* (Dd). (**E**) T_m_ predictions of *I. cascadensis* compared to *V. vermiformis*. (**F**) Predicted changes in net surface charge, fraction of positively charged surface residues, and fraction of negatively charged residues between the protein sequences of *I. cascadensis* and *V. vermiformis*. Dots are analogous proteins, and each pair is connected by a line. Red and blue lines indicate a corresponding decrease or increase from *I. cascadensis* to *V. vermiformis*. (**G**) The surface of dihydropteridine reductase from *I. cascadensis* and *V. vermiformis*. Red and blue regions indicate negative and positive electrostatic surface potential, respectively.

We profiled the gene content of *I. cascadensis* and three reference Amoebozoa genomes (**Fig. 4A; Table S9**). Comparison from orthogroup clustering across the genomes revealed 97% of orthogroups shared between the two *I. cascadensis* assemblies, indicating robustness of independent *de novo* assemblies (**Fig. 4A**). *I. cascadensis* and *V. vermiformis* shared 52% of orthogroups with each other and less than 40% with the other amoebae. We identified putative functions and pathways enriched in *I. cascadensis* relative to more mesophilic amoebae (*41*). Signaling networks and pathways including calcium, mitogen-activated protein kinase (MAPK), and Ras, were all enriched in *I. cascadensis* (FDR adj. P<0.0001 for all; **Fig. 4C**; **Table S10**). Calcium signaling pathways can initiate protective responses to thermal fluctuations by altering lipid metabolism (*42*, *43*). The rapid signaling and initiation of cellular responses to heat is consistent with the ecology of this amoeba, which lives in a dynamic geothermal stream environment.

*Incendiamoeba cascadensis* may also employ an expanded molecular toolkit to manage proteostasis. Genes hypothesized to encode molecular chaperones including heat shock proteins (HSPs; HSP20/small HSPs and HSPA5) were enriched in *I. cascadensis* (**Fig. 4D**; **Table S10**) and transcribed in *I. cascadensis* transcriptomes. Small HSPs prevent protein aggregation and assist in protein refolding, while HSPA5 regulates the unfolded protein response (UPR) (*44*). Genes encoding a suite of ubiquitin-protein ligases, which aid in protein degradation (*45*) including RLIM (*46*) and HERC2, involved in DNA repair regulation (*47*), were also enriched and transcribed (**Fig. 4D**).

In addition to characterizing gene content, we used the complete set of predicted protein sequences from *I. cascadensis* and *V. vermiformis* to obtain predicted melting temperature (T_m_) distributions, using TemBERTure to capture overall trends (*48*, *49*). The T_m_ is the temperature at which the population of a protein is 50% unfolded and is commonly used to quantify protein thermal stability. We emphasize that this prediction is not accurate for absolute melting temperatures of individual proteins but can capture trends in overall distributions (*48*). With this in mind, the mean predicted melting temperature of the *I. cascadensis* proteome is significantly higher, 50.9 ± 6.6°C, in comparison to 46.9 ± 5.1°C for *V. vermiformis* (P < 0.0001; **Fig. 4E**). *I. cascadensis* is also predicted to have five times more proteins with T_m_ over 60°C than *V. vermiformis*, which may reflect increased protein stability. This elevated T_m_ for a thermophile is in line with other studies comparing thermophilic and mesophilic organisms (*49*, *50*).

*I. cascadensis* proteins exhibit distinct biophysical properties that likely contribute to their stability under heat stress. We predicted protein structure from 30 highly transcribed sequences in *I. cascadensis* and *V. vermiformis* using Alphafold2 (*51*, *52*) and compared a range of structure-derived features (**Table S11**, **Fig. 4F**, **Fig. S7**). Our analysis showed that *I. cascadensis* proteins displayed a significantly higher net surface charge (FDR adj. P = 0.002), reflecting an enrichment in the fraction of positively charged residues (FDR adj. P = 0.005) and a depletion in fraction of negatively charged residues (FDR adj. P = 0.005) on the protein surface. These differences in surface charge, (for example, visualized for surface of the protein dihydropteridine reductase, a critical enzyme in the folate biosynthesis pathway (*53*); **Fig. 4G**), may enhance protein stability at elevated temperatures by promoting favorable electrostatic interactions, consistent with prior efforts linking surface charge to thermal stability (*54*, *55*).

### Conclusions

*Incendiamoeba cascadensis* proliferates at temperatures beyond what was thought possible for any eukaryotic organism. This discovery raises new questions about the true maximum temperature a eukaryotic cell can endure. While red algae and fungi are generally considered to host the most thermophilic eukaryotic lineages (*12–16*), we show that amoebae are particularly successful in high-temperature environments, and our biogeographic analysis indicates that additional thermophilic species within the clade await discovery. Our findings suggest molecular and ecological strategies for thermostasis in eukaryotes and provide an untapped source of thermostable proteins with promising biotechnological applications, from industrial enzymes to novel protein stabilization mechanisms for synthetic biology. Likewise, these results have profound implications for our understanding of the evolutionary constraints on eukaryotic cells and the set of abiotic parameters that inform the search for life elsewhere in the Universe.

## Acknowledgments

We acknowledge QB3 Genomics, UC Berkeley, Berkeley, CA, RRID:SCR_022170 for sequencing and thank Eva Myšková for assistance with library preparation. We thank Danielle Jorgens and Reena Zalpuri from the University of California Berkeley Electron Microscope Laboratory for advice and assistance in TEM sample processing and imaging. We also thank Arthur Charles-Orszag for coordination and introduction to the heated microscope setups. Thank you to David Booth and his lab members for sharing live cell reagents. FM thanks the EMBL Advanced Light Microscopy Facility for equipment and technical expertise. The work conducted by the U.S. Department of Energy Joint Genome Institute (https://ror.org/04xm1d337), a DOE Office of Science User Facility, is supported by the Office of Science of the U.S. Department of Energy operated under Contract No. DE-AC02-05CH11231.

## Funding

National Science Foundation Graduate Research Fellowship (HBR)

National Science Foundation Division of Environmental Biology Award 2439029 (AMO)

National Aeronautics and Space Administration Exobiology Program grant 24-EXO24-0042 (AMO)

American Philosophical Society Lewis and Clark Fund for Exploration and Field Research in Astrobiology (AMO)

National Institute of General Medical Sciences of the National Institutes of Health Award R35-GM118119 (RDM)

Howard Hughes Medical Institute Investigator program (RDM)

U.S. National Science Foundation Division of Environmental Biology Awards 243903 and 2230391 (LAK)

U.S. National Science Foundation Division of Molecular and Cell Biology Award 2025305 (KMS)

National Institute of General Medical Sciences Award 1R16GM154184 (KMS) European Commission KaryodynEVO, 101078291 (GD)

European Molecular Biology Laboratory (GD)

Gordon and Betty Moore Foundation GBMF13113 (GD, FM, and OD) Swiss National Science Foundation Starting Grant TMSGI3_218007 (OD)

## Author contributions

Conceptualization: HBR, AMO

Field sampling: HBR, AMO, KMS, IH, GVW, TT, KS

Culturing: HBR, RMS, TT, NAP, AMO

Expansion microscopy: FM, NAP, OM, GD

SEM imaging: NAP

Heated microscopy: HBR, NAP, KS, SJL, RDM

Motility analyses: KS, EM, RDM

Phylogenomics: GA, LAK

18S phylogenetic tree construction: TT

Protein analysis: SS, JKN, HBR

Bioinformatics and genomics: KL, HBR, FS, AMO

Writing: HBR, AMO with contributions from all coauthors.

## Data and materials availability

Raw reads and assemblies for all genomes generated in this study have been deposited in the NCBI Sequence Read Archive in BioProject PRJNA1329177. 18S rRNA gene sequences of *I. cascadensis* are deposited in NCBI for strain HSC4 (LC894571), HSC6 (LC894572), and type strain NS2 (LC894573). The 18S sequence of *V. vermiformis* strain HSC3 is deposited in NCBI under LC894570. Code and data files are available on Github (https://github.com/hbrappaport/incendiamoeba_manuscript), and an archived version is available from the following Figshare dois: 10.6084/m9.figshare.30686090, 10.6084/m9.figshare.30686081, 10.6084/m9.figshare.30686078, 10.6084/m9.figshare.30686072, 10.6084/m9.figshare.30686063, 10.6084/m9.figshare.30685748

## Supplementary Materials

Materials and Methods

Supplementary Text

Figs. S1-S8

Tables S1-S12

References (56–104)

Movies S1-S12

