## Supplementary Materials for "A geothermal amoeba sets a new upper temperature limit for eukaryotes"

Materials and Methods

Supplementary Text

Figs. S1-S8

References (56-104)

Tables S1-S12

Movies S1-S12

Materials and Methods

Field sampling

We collected sediment, microbial biofilm, and water samples from 11 sites along a tributary of Hot Springs Creek (40º 26.510’N, 121º 24.167W, ~84m transect), located in the Warner Valley in Lassen Volcanic National Park, CA, USA in August 2023, July 2024, and July 2025 under permits LAVO-2023-SCI-0028, LAVO-2024-SCI-0017, and LAVO-2025-SCI-0016. Although most Lassen geothermal features are acid-sulfate steam-heated, the tributary is a part of Drakesbad, a low chloride, neutral pH hydrothermal system that is likely heated from a shallow source (*20*). Sampling site temperature ranged from 47-64°C and pH averaged 7.55, measured in the lab with a pHenomenal pH meter (VWR; **Table S1**). We collected and transported samples in sterile 50mL conicals and 1L polycarbonate bottles and kept them at room temperature until cultivation.

Cultivation and isolation

To increase the density of protists from the samples, we established initial enrichment cultures in 50mL plastic tissue culture flasks with 1mL of sample and 19mL of autoclaved Volvic^®^ water within a few days of sampling (*17*). We added sterile wheatberries as a nutrient source to grow bacteria sourced from the same environment, which serve as prey for the target protists (*56*). We kept flasks in the dark, so although we observed eukaryotic algae and cyanobacteria in initial cultures, no photosynthetic organisms remained in enrichments. We incubated enrichments across a temperature range from 37°C to 60°C. *Incendiamoeba cascadensis* became dense and emerged as the sole eukaryotes in cultures incubated at ≥ 55°C. We maintained cultures in 250mL flasks with Page’s Amoeba Saline (*57*) and transferred cultures into fresh media weekly (*58*, *59*).

Growth experiments

To determine the growth range of the novel species, we conducted growth experiments in a manner similar to (*17*). First, we grew amoeba cultures for at least a week to high density and then measured cell density in four replicates with a Neubauer hemocytometer (NanoEntek C-chip 4 chamber). We pipetted dense amoeba stock into 50mL tissue culture flasks with a single autoclaved wheatberry and Page’s amoeba saline to reach a final density of 2.50E+04 cells/mL in 10mL. We set up four replicate flasks per temperature and confirmed initial density with a hemocytometer before incubating the flasks at a range of temperatures from 30°C to 64°C. Every 24 hours, we removed flasks from the incubators, imaged them, filled to 10mL with autoclaved DI water to retain a stable salinity after evaporation, and we allowed amoebae to detach from the flask surface at room temperature. After 30 minutes, we gently shook flasks and counted each replicate once with the hemocytometer. After seven days, we calculated the average doubling time for each temperature for the exponential growth period (**Table S5**). Doubling times were variable, as factors such as initial bacterial input and evaporation likely impacted cell growth in addition to temperature. We cross-checked incubator temperatures with EasyLog Temperature Data Loggers (Lascar Electronics) recording temperature every ten minutes (**Table S4**) and a NIST-certified Traceable Datalogger. We used temperature experiments to determine the upper and lower boundaries of cellular replication and to estimate the optimal growth temperature.

Temperature-controlled microscopy

We imaged *I. cascadensis* in a Bioptechs Delta-T (*33*) dish or the Interherence VaHeat heated stage. We washed Bioptechs dishes with MilliQ water then primed with sterile-filtered, conditioned growth media. We calibrated dish temperature with a Fisherbrand Traceable Big-Digit Type K Thermometer (0.1ºC resolution, NIST certified 1ºC precision). We then replaced conditioned media with 2.5mL of dense, freshly detached culture, brought the dish to the appropriate temperature, and then imaged. We plasma cleaned VaHeat substrates, coated them with Poly-L-Lysine (Sigma P8920), and washed with fresh Page’s Saline. We then treated substrates with sterile-filtered conditioned growth media for at least 30 minutes prior to exchange with freshly detached cells.

For morphological characterization, we performed live cell imaging at 100X as previously described in (*33*) with the exception that the objective was heated to 57ºC for imaging at 60ºC and heatshock experiments, and 60ºC for imaging at 63ºC. We performed reflective interference contrast microscopy (RICM) using a Nikon TI2-LA-FL-3 epi illuminator on a Ti2 microscope, fitted with a liquid light guide illuminated by the Cy3 channel of a Lumencor SpectraX LED source. The dichroic cube held a Chroma 50/50 beamsplitter with a D546/10X exciter filter. We adjusted illumination intensity and the aperture diaphragm on the epi illuminator to achieve the optimal contrast. Dark regions of the RICM image correspond to areas of close contact between the cell membrane and the glass coverslip. We imaged Differential Interference Contrast (DIC) and RICM at 100X using a Photometrics BSI camera. To describe nuclear morphology, we stained nuclei with two drops of NucBlue (ThermoFisher) per dish and excited samples with a 405 laser. We stained actin with 1µM SiR-Actin (Cytoskeleton).

For motility analysis, we imaged cells by DIC using a 40X Nikon Plan Apo (NA 0.95) objective on a Nikon Ti-E microscope fitted with a Chameleon3 series monochrome camera (Point Grey Research/Teledyne) and controlled by Micromanager 2.0.3. We acquired 300 frames at one frame/second using one field of view.

SEM and TEM sample preparation

To fix *I. cascadensis* for SEM, we detached cells from culture flasks grown at 58 or 63ºC by pouring off media (~60mL) and resuspending in 5-7mL room temperature Page’s Saline. We gave cells ten minutes to detach and then plated on plasma cleaned and Poly-L-Lysine (Sigma P8920) coated 12mm high precision coverslips (Thorlabs CG15NH) in a 24-well plate. We returned plates to the 58ºC incubator for 30-60 minutes. We fixed cells by adding an equal volume of prewarmed 2x fixation solution (final concentration 1X PHEM buffer and 4%PFA or 1X PHEM buffer and 2% glutaraldehyde) for 30 minutes, then washed with 0.1M HEPES pH 7.0 and stored at 4ºC. To fix *V.vermiformis* for SEM, we transferred plasma cleaned and Poly-L-Lysine coated coverslips to a 6-well plate (multiple coverslips per well). We split cells into the 6-well plate using fresh Page’s Saline and a wheatberry and allowed cells to grow for one to three days. We fixed cells at the highest density of amoebae by adding 2x fixation solution (final concentration 1X PHEM buffer and 4% PFA). For heat shock, we transferred settled *I. cascadensis* cells to a 70ºC incubator and added 2x fixation buffer (final concentration 1X PHEM and 4% PFA) at the appropriate time points.

For SEM, we performed secondary fixation with 1% osmium tetroxide for one hour at room temperature in the dark. Then, we dehydrated cells with 25%, 50%, 75%, 90%, 100% ethanol (the ethanol for the last step had been dehydrated using 4Å, 8-12 mesh molecular sieves that had been rinsed and baked) and critical point dried. We then sputter coated samples with 5nm iridium. We performed SEM imaging on a Phenom Pharos tabletop FEG-SEM.

To prepare samples for TEM, we seeded amoeba cultures grown to high density onto agar plates (2.5% BactoAgar in Page’s saline) and incubated at 57°C for one hour. We fixed adhered amoeba trophozoites *in situ* with a mixture of 2.5% paraformaldehyde and 2.5% glutaraldehyde in 0.1M sodium cacodylate buffer for 30 minutes, postfixed with 1% osmium tetroxide and 1.6% Potassium ferricyanide for two hours, dehydrated with a series of acetones, and embedded in Epon resin. We examined ultrathin sections stained with 2% aqueous uranyl acetate and 0.3% lead citrate 0.1 N sodium hydroxide using the Tecnai 12 electron microscope at the University of California Berkeley Electron Microscope Laboratory.

Ultrastructure expansion microscopy (U-ExM)

We performed U-ExM as previously described with slight modifications (*60*). We grew cells on plasma cleaned and Poly-L-Lysine coated coverslips at 58 or 63ºC and fixed in 1X PHEM buffer with 4% paraformaldehyde for 30 minutes. Next, we carefully washed coverslips with 0.1M HEPES pH 7.0 and shipped them in PBS. For U-ExM, we anchored coverslips overnight at 37ºC in 0.7% formaldehyde and 1% acrylamide, embedded them into a swellable hydrogel, and polymerized for 45 minutes at 37ºC in a humid chamber. We denatured samples at 95ºC in an SDS containing buffer for 1.5 hours, washed with PBS twice, and expanded in three successive incubations in ddH2O for 15 minutes each. We measured gel diameters for use as a proxy for cell expansion factors.
We stained after shrinking the gels with three PBS washes. For antibodies, we used ABCD (AntiBodies Chemically Defined; Geneva Antibody Facility), antibodies against alpha- and beta-tubulin (AA345 and AA344) and (AF291) at 1:500 in 3% BSA in PBS-T (0.1 % TritonX-100) and incubated at 37ºC overnight. We removed the unbound antibody by three PBS-T washes for five minutes each before incubating gels in secondary antibodies, anti-mouse AlexaFluor488 (Thermo A-11001) and anti-guineapig AlexaFluor647 (Thermo A-21450), for four hours at 37ºC. We again washed gels three times with PBS before staining with Bodipy-TR ceramide (Invitrogen D7540) at 1:2000 and Hoechst 33342 (ThermoFisher 62249) for three hours at 37ºC, washing again with PBS and re-expanding them in ddH2O for imaging.
We acquired images with a Nikon-CSU-W1 SORA 750 spinning disk microscope with a SR P-Apochromat IR AC 60X WI/1.27 objective. We mounted gels onto PLL coated Ibidi glass bottom chambers for imaging to reduce sample drift.

Image analyses

We measured cell size from 2,414 cells across ten temperature conditions (**Table S3**). We measured nuclear diameter from 29 cells imaged at 60ºC. To remove uneven illumination and background artifacts from high temperature, we first processed image sequences using a combined fast Fourier transform and bandpass filter in FIJI (Version 2.16.0/1.54p, (*61*)). We filtered large structures down to 50 pixels, small structures up to five pixels, with no stripe suppression and 5% tolerance of direction. To identify the cell edge for cell shape and tracking, we first duplicated each image sequence. We inverted one image and thresholded both to generate binary masks. We then combined the masks to include both the illuminated cell edges as well as the shadows from differential interference contrast.

Once images were segmented, we conducted particle analysis in FIJI with the following conditions: circularity=0.1-1.00 and size >75µm. We excluded particles on edges to restrict analysis to whole cells and optimize downstream tracking. Size thresholding removes non-filamentous bacteria from the analysis, and circularity thresholding removes the majority of filamentous bacteria. We recorded all thresholded cells as outlines. We extracted major and minor axes, area, and centroids along with frame number using a custom macro, and we recorded each measurement per frame. In FIJI, we determined centroids by averaging all xy coordinates in the thresholded area and defined major and minor axes by fitting an ellipse to the object and identifying its primary and secondary axes (Fiji Documentation: https://imagejdocu.list.lu/start). We defined aspect ratio as the ratio between the major and minor axes. We reported axes and aspect ratio measurements as averages (**Table S3, Table S6**) or per frame (**Fig. 3D**).

To track cell positions, we assigned particle IDs using a custom script in R (R version 4.3.2). We used the k-nearest neighbors algorithm through the "Fast Nearest Neighbors" R library to link particles across frames. We set maximum allowed displacement per particle between frames and gap tolerance criteria, or number of frames a particle can disappear and still retain its ID, during each run. Maximum allowed displacement per particle filters completely detached cells. We removed particles appearing in five or fewer frames and assessed performance of the particle-linking process by visual track display (**Fig. S5A**) and by particle ID assignment integrity post hoc. We used track length (**Fig. S5B**) as a metric for quality control. As fast-moving particles tended to move out of field of view, we performed correlation between track length with measured variables to check for bias introduced by tracking quality. We found no significant correlations between any measured metric and track length (**Fig. S5B**).

 	We used a custom R script to quantify the directional change of particles during movement. Briefly, we calculated displacement vectors, movement angles, and angle changes over time for individually tracked particles. We summarized the mean absolute angle change per track and normalized to the total time for the tracked particle.

We also used a custom R script to calculate the mean squared displacement (MSD) of particles over time. Briefly, we dynamically set the maximum time lag (τ) for displacement calculations based on the median track length for each condition, minimizing bias from short particle tracks. We calculated squared displacement as Δx^2^+Δy^2^ for x,y particle positions. The MSD scales with time lag (MSD∝τ^α^) where we determined α from the slope of a log-transformed plot of MSD versus τ (**Table S7**). To understand the type of motion, we statistically compared α against a reference value of 1 using two one-sided t-tests. Values greater than 1 indicate directed motion. We used tracked particles with trajectories filtered by maximum time lag (τ) to calculate the convex hull area using the chull function in R. Convex hull is the minimal convex polygon that contains the particle trajectory. We then used the shoelace formula for polygons to compute the area from extracted coordinates to quantify extent of space exploration.

Molecular cloning and phylogenetic analysis

We extracted DNA from established monoeukaryotic cultures using the Monarch Spin gDNA Extraction Kit (New England Biolabs). We amplified nearly the entire 18S rRNA gene sequence of *I. cascadensis* (strain NS2, 2023) and *V. vermiformis* (strain HSC3, 2024) by PCR using universal eukaryotic primers EukA and EukB (*62*). PCR conditions followed the Q5 Hot Start Master Mix protocol (New England Biolabs): 98ºC for 30 seconds, 34 cycles of 98ºC for 10 seconds, 67ºC for 30 seconds, 72ºC for 1 minute, then 72ºC for 2 minutes. We visualized the resulting amplicons by gel electrophoresis and then excised and purified bands of approximately 3 kb (*I. cascadensis*) and 1.8 kb (*V. vermiformis*) using the Monarch DNA Gel Extraction Kit (New England Biolabs) according to the manufacturer's instructions. Using the TOPO TA Cloning Kit (Invitrogen) and following the manufacturer's protocol for chemical transformation, we ligated purified amplicons into the pCR4-TOPO vector and transformed into One Shot competent cells, plated on LB agar with kanamycin (50 µg/mL), and screened by PCR using M13 primers. We extracted plasmid DNA from overnight cultures using the Monarch Plasmid Miniprep Kit (New England Biolabs) according to the manufacturer's instructions. We sequenced at the UC Berkeley Sequencing Center using M13 primers and internal primers NS2_975F (5′-TCGTTAACGGAATTAACCAGAC-3′) and 1350R (*63*) for the longer NS2 inserts.

We inspected the 18S rRNA gene sequences for ambiguous base calls and assembled the sequences in Geneious Prime v2025.0.3 (Biomatters) before subjecting them to BLAST (*64*) searches against the GenBank (NCBI) and IMG/M (*65*) databases to identify top hits. To search for previous detection in IMG/M metagenomes, we queried the *I. cascadensis* 18S rRNA gene sequence against the “18S rRNA public assembled metagenomes 2025-06-16” JGI IMG database (https://img.jgi.doe.gov) containing 2,509,354 sequences extracted from 31,093 publicly available genomes. To infer the phylogenetic positions of *I. cascadensis* and *V. vermiformis* strain HSC3 with 18S gene sequences, we compiled a sequence dataset comprising the top BLAST hits and representatives of major Tubulinea lineages with Centramoebidae as the outgroup. We aligned the dataset using MAFFT v7 (*66*) with the Q-INS-i algorithm. We trimmed the alignment using trimAl v.12 (*67*) with trimming thresholds -gt 0.3 -st 0.001, resulting in a final alignment of 1,437 bps, and we inspected the alignment in Jalview v.2 (*68*). We computed the final tree with the IQ-TREE web server (*69*) with TIM3+F+R7 model selected by ModelFinder with Bayesian information criterion (*70*). We tested tree branches by the standard bootstrap analysis (*71*) and the approximate Bayes test (*72*). We visualized the tree in iToL (*73*).

Genome and transcriptome sequencing

To target amoeba genomes and transcriptomes, we concentrated two sets of four 60mL cultures by centrifuging for ten minutes at 1000g. We added 200µL DNA/RNA shield (Zymo) to the pellet and extracted DNA with the QuickDNA HMW MagBead Kit (Zymo) according to the protocol for cultured cells. We extracted RNA with the Monarch Total RNA Miniprep Kit (New England Biolabs) according to the protocol for mammalian whole blood. We prepared ultra-low input DNA with the SMRTbell Express Template Prep Kit 2.0 (PacBio) and sequenced in a pooled library with PacBio Revio 25M SMRT Cell at QB3 Genomics, UC Berkeley, resulting in 6.6GB of data and 3,570,574 sequencing reads. We separated the amoeba from bacterial or viral contigs, resulting in a single *I. cascadensis* bin from each sample (**Fig. 4A**). We deduplicated reads using smrtlink_13.0.0.207600.sif pbmarkdup. For read quality control, we used BBDuk (version 39.19; (*74*)) to remove reads that contained PacBio control sequences and PCR adapters and used IceCreamFinder (version 39.19; (*74*)) to remove reads without SMRTbell sequences. We then assembled reads with Flye (v2.9.5-b1801; (*75*)) and binned with MetaBAT2 (v2.15-5-g1a9bac2; (*76*)). To assess bin completeness and contamination, we assessed presence or duplication of Benchmarking Universal Single-Copy Orthologs (BUSCOs; (*39*)), identifying the single largest bin with highly complete eukaryotic markers as the *Incendiamoeba* draft assembly. We estimated additional completeness in light of shared missing markers across Amoebozoa (as the BUSCO marker set is not specific to Amoebozoa) by adding shared missing markers to our estimate. We screened for a mitochondrial genome using an in-house tool (screen_mito2). We predicted genes from the assembly using Prodigal (*77*) and searched against an in-house curated database of mitochondrial HMMs using HMMER hmmsearch version 3.1b2 [--domtblout] (*78*) and compared these results with predictions from MFannot (*79*).

To obtain metagenomes for characterization of the enriched bacterial community, we filtered ~60mL of four dense cultures onto 0.2µm filters and extracted DNA with the DNEasy PowerSoil Kit (Qiagen) following manufacturer’s instructions. We sent extracted DNA to SeqCoast Genomics (Portsmouth, NH) for Illumina sequencing, generating 13.3 million 150bp paired end reads. We filtered bacterial metagenome assembled genomes through KBase with Trimmomatic (*80*), assembled with metaSPAdes v3.15.3 (*81*), and binned with MaxBin2 v2.2.4 (*82*), CONCOCT v1.1 (*83*), and MetaBAT2 v1.7 (*76*). We optimized bins with DasTool v1.1.2 (*84*). We assessed the quality of extracted bins with CheckM v1.0.18 (*85*) and assigned taxonomy with GTDB-Tk v2.3.2 (*86*). The 17 genome bins assembled collectively comprise >85% of the total reads, indicating the dominant bacterial community is well recovered at the genome level. We constructed a phylogenetic tree from these bins using KBase SpeciesTree v2.2.0, which relies on a set of 49 core, universal genes defined by COG gene families.

We PolyA selected and prepared transcriptomes with the Nextera XT kit and sequenced 150bp reads with Illumina NovaSeq X 10B. To remove bacterial sequences, we concatenated the 17 bacterial MAGs assembled from metagenomes and mapped transcripts against the resulting database using BBMap (version 39.33; (*87*)). We then mapped the unmapped reads to *I. cascadensis* assembly. We predicted gene calls with BRAKER3 (*88*) from RNA reads and draft genomes. We then tabulated paired-end reads assigned to each BRAKER gene (e.g. read counts) with featureCounts (*89*). We merged transcript counts with KEGG orthology assignments and annotations. We filtered read counts to retain genes which had a minimum total count of ≥10.

Phylogenomics and functional analyses

We input BRAKER3 gene calls to EukPhylo (*23*) to produce gene trees for 596 gene families conserved across Amoebozoa (71 Amoebozoa taxa and 253 total taxa). Next, we produced a phylogeny from a multi-sequence alignment using 102 gene families selected from an initial set of 270 based on a minimum of 30 Amoebozoa taxa and both novel amoeba replicates after running gene calls through EukPhylo. To assess the likelihood that the novel amoeba belonged in the same genus as its nearest neighbor, *Vermamoba vermiformis*, we computed the average distance separating other species that fall within the same genus in our Amoebozoa phylogeny (**Fig. 1A**). To do this, we calculated the branch length to the nearest shared common ancestor data from six within-genus pairs of species from the following genera: *Nebela*, *Hyalosphenia*, *Difflugia, Acanthamoeba, Paramoeba,* and *Dictyostelium* (**Table S2**). Then, we assessed whether the distance between the novel amoeba and *V. vermiformis* fell within a 95% confidence interval from the known within-genus pairs of species across the phylogeny.

To elucidate which gene functions and pathways may be present or enriched in *I. cascadensis* relative to other amoebae, we obtained publicly available genomes from three additional species of Amoebozoa. We ran these three genomes through BRAKER3 to facilitate comparisons (*Vermamoeba vermiformis* CDC-19 GCA_045837725.1 (*90*), *Acanthamoeba terricola* Neff GCA_000313135.1 [previously *castellanii*] (*91*), and *Dictyostelium discoideum* GCA_000004695.1 (*92*)). We inferred the orthogroups of the two *I. cascadensis* draft genomes and three comparative Amoebozoa genomes using OrthoFinder (*93*). We calculated the proportions of shared orthogroups between genomes from means of the OrthoFinder output pairwise comparisons of total orthogroups. We functionally classified predicted genes using the KEGG Automatic Annotation Server with the bi-directional best-hit method. Following the metric used in (*41*), we calculated enrichment by adding the number of species that had fewer orthologues than *I. cascadensis* and calculated depletion by subtracting the number of species that had more orthologues. Then, we used a non-parametric, permutation-based approach to assess whether the number of enriched or depleted gene functions for each pathway/category (BRITE level C) significantly differed from a random sampling. Briefly, we performed 10,000 random permutations of enrichment types (enriched, depleted, neutral; where < -2.5 = enriched, ≥ -2.5 ≤ 2.5 = neutral, and >2.5 = depleted) to simulate a random distribution and then compared observed counts to our permuted counts (two-sided p-values were FDR adjusted for multiple tests).

Protein Analyses

To compare predicted melting temperature (T_m_) distributions of *I. cascadensis* and *V. vermiformi*s, we used the complete set of predicted protein sequences. We obtained T_m_ predictions with TemBERTure, a deep learning framework trained on 48,000 protein melting temperatures (*48*, *49*). We calculated summary statistics for both species' sets of protein sequences and compared T_m_ means with a one-sided Welch’s t-test. To analyze biophysical properties, we predicted the structure of 30 of the most abundant proteins involved in core cellular processes using AlphaFold2 through ColabFold v1.5.5 with standard parameters, reasoning that widespread unfolding of these proteins will lead to highly deleterious effects on the cell (*51*, *52*, *94*). We then analyzed these PDB structures using MDTraj (*95*), MDAnalysis (*96*, *97*) and OpenMM (*98*). Analyses included radius of gyration, secondary structure composition using DSSP, solvent-accessible surface area with the Shrake-Rupley algorithm, and intramolecular interaction quantification (**Table S11)**. We performed statistical comparisons between the calculated metrics of *I. cascadensis* and *V. vermiformis* proteins using a two-sided dependent T-test, with a Benjamini-Hochberg correction for false discovery (**Fig. 4, Fig. S7**). The electrostatic surface potential of the protein dihydropteridine reductase was calculated using Adaptive Poisson-Boltzmann Solver (APBS) (*99*) and visualized using PyMOL (*100*)**.**

Supplementary Text

Taxonomic appendix

Amoebozoa Luhe 1903, sensu Cavalier-Smith 1998

• Tubulinea Smirnov et al. 2005

•• Echinamoebida Cavalier-Smith, 2004, Smirnov et al, 2011

••• Vermamoebidae Cavalier-Smith and Smirnov 2011

***Incendiamoeba* gen. nov.** Rappaport, Petek-Seoane, Tyml, Mikus, LaButti, Ani, Niblo_,_ MacVicar, Shepherd, de la Higuera, Lord, Dey, Wolfe, Dudin, Sukenik, Katz, Stedman, Skruber, Schulz, Mullins, et Oliverio

**Diagnosis:** Lobose amoeba; trophozoites limax-shaped with prominent anterior hyaline cap in directional locomotion; highly polymorphic in non-directed movement, rapidly shifting between polytactic and flattened amoebiform states; movement with occasional eruptive activity; cysts double-walled; typically mononuclear with complex nucleus; mitochondria with non-branching tubular cristae; cell surface with amorphous glycocalyx.

**Etymology:** Incendiamoeba (In.cen.di.a.mo’e.ba. N.L. fem. n. *Incendiamoeba*, composed of L. n. *incendium*, fire, conflagration, and N.L. fem. n. *amoeba*; “fire amoeba” referring to the thermophilic nature of the organism)

**Type species:** *Incendiamoeba cascadensis*

***Incendiamoeba cascadensis* sp. nov.** Rappaport, Petek-Seoane, Tyml, Mikus, LaButti, Ani, Niblo_,_ MacVicar, Shepherd, de la Higuera, Lord, Dey, Wolfe, Dudin, Sukenik, Katz, Stedman, Skruber, Schulz, Mullins, et Oliverio

**Diagnosis:** As for the genus described above; trophozoites in directional locomotion (vermiform): length 38.7 ± 9.4 μm, width 10.5 ± 3.1 μm, length:width ratio 3.9 ± 1.2 (n = 135); amoebiform state: length 23.5 ± 7.9 µm, width 13.0 ± 3.5 µm, length:width ratio 1.8 ± 0.4 (n = 1176); nucleus spherical, averaging 4.0 μm in diameter (n = 29). Values are reported as mean ± standard deviation for tracked cells.

**Etymology**: *cascadensis* (cas.ca.den'sis. N.L. fem. adj. *cascadensis*, from the Cascades, referring to the Cascade Range of western North America, hosting the geothermal site where the amoeba was isolated)

**Type Location:** Lassen Volcanic National Park, California, United States (40° 26.510' N, 121° 24.167' W). Geothermal spring sediment, collected near Hot Springs Creek, a neutral pH and high temperature (65ºC at 40º 26.497’N, 121º 24.140’ W to 35ºC at 40º 25.533’ N, 121º 24.177’ W) creek within the Drakesbad area.

**Type material:** Type culture will be deposited with the Culture Collection of Algae and Protozoa (UK) and the German Collection of Microorganisms and Cell Cultures. The sequence of the 18S rRNA gene is deposited with Genbank under the accession number LC894573.

**Remarks:** We took all measurements in Bioptechs glass-bottom heated dishes at 57-60ºC.

Additional evidence for optimal motility range

To further characterize movement, we analyzed directional changes between successive positions to understand when cells were conducting smooth and persistent versus oscillatory motion. Angle changes centered at 0º on plotted histograms (**Fig. S6B**) indicate more unidirectional motion, while changes centered around ± 180 indicate frequent reversal. Histograms show that directed motion occurred most frequently from 57-64ºC. While cells at 55ºC still exhibited some directional motion, as temperature rose to 66ºC or decreased to 44ºC, motion became predominantly oscillatory. In contrast, motion was highly directional for the species *V. vermiformis* at 25ºC (**Fig. S6B**). By inscribing a cell's trajectory within a polygon, its spatial footprint can describe the amount of space explored. Between 57-62ºC the spatial footprint (determined from the convex hull area, **Supplementary Methods**) averaged 34.1µm² while cells at 44 and 66ºC explored areas of 5.5µm² and 1.8µm², respectively (**Table S7**). Unlike metazoa, amoeba likely lack specific adhesion complexes for surface attachment (*101*). Cell area provides insight into the extent of substrate engagement and is a proxy for adhesion, which is required for force transmission to the surface during motility (*102*, *103*). We recorded an over two-fold increase in cell area from encystment temperatures (25ºC, 66ºC, 70ºC) to amoebae with the highest motility rates and spatial exploration (57-62ºC, **Fig. S6A**, **Table S7**).

Metabolically diverse bacterial consortium grows with *Incendiamoeba*

To characterize the bacteria in culture with *I. cascadensis,* we sequenced the strain NS2 metagenome. We recovered a total of 17 bacterial species-level genomes from metagenomic sequencing data (**Fig. S8A**), 11 of which represent putatively novel species based on average nucleotide identity (ANI) and relative evolutionary divergence (RED values reported from GTDB-Tk; **Fig. S8; Table S12**). Notably, *Meiothermus ruber*, a globally distributed red-pigmented, filamentous Deinococcota bacterium found in natural and artificial thermal environments (*28*), made up 41.34% of all reads (**Fig. S8B**). The growth range of the *M. ruber* type strain is 35–70°C, with an optimum temperature of 60°C (*28*), aligning with the growth range and optimum of the novel amoeba. Based on feeding of filamentous bacteria (**Movie S5; Fig. S3**) and relative abundance (**Fig. S8B**), we hypothesize that *M. ruber* is the primary food source of the novel amoeba, acknowledging that the enriched bacterial community grown in the dark likely differs in composition from the *in-situ* community, which may offer a suite of other prey sources. Other consortium members include *Pelomicrobium methylotrophicum*, isolated from a mud volcano (*104*), species of *Chloroflexia* and *Thermoleophilum*, both known thermophilic groups, and novel species within Terriglobia, Kapabacteria, and *Caldilinea.*

We predict the bacterial community in these LVNP cultures to be metabolically diverse (**Fig. S8C**). Along with variation in aerobic respiration pathway genes among consortium members, there was high variation in genes for carbohydrate active enzymes (CAZy) such as cellulose and glucans, as well as nitrogen, methane, and sulfur metabolism, arsenate and mercury reduction, and alcohol production (**Fig. S8C**), suggesting syntrophic interactions involving the exchange of metabolic byproducts, such as organic acids, alcohols, or gases, between members to sustain collective metabolic processes. This may suggest why *I. cascadensis* has not been previously discovered, as much of the previous culturing of geothermal amoebae has relied on isolating a single bacterial strain as a food source rather than community enrichments (*17*).

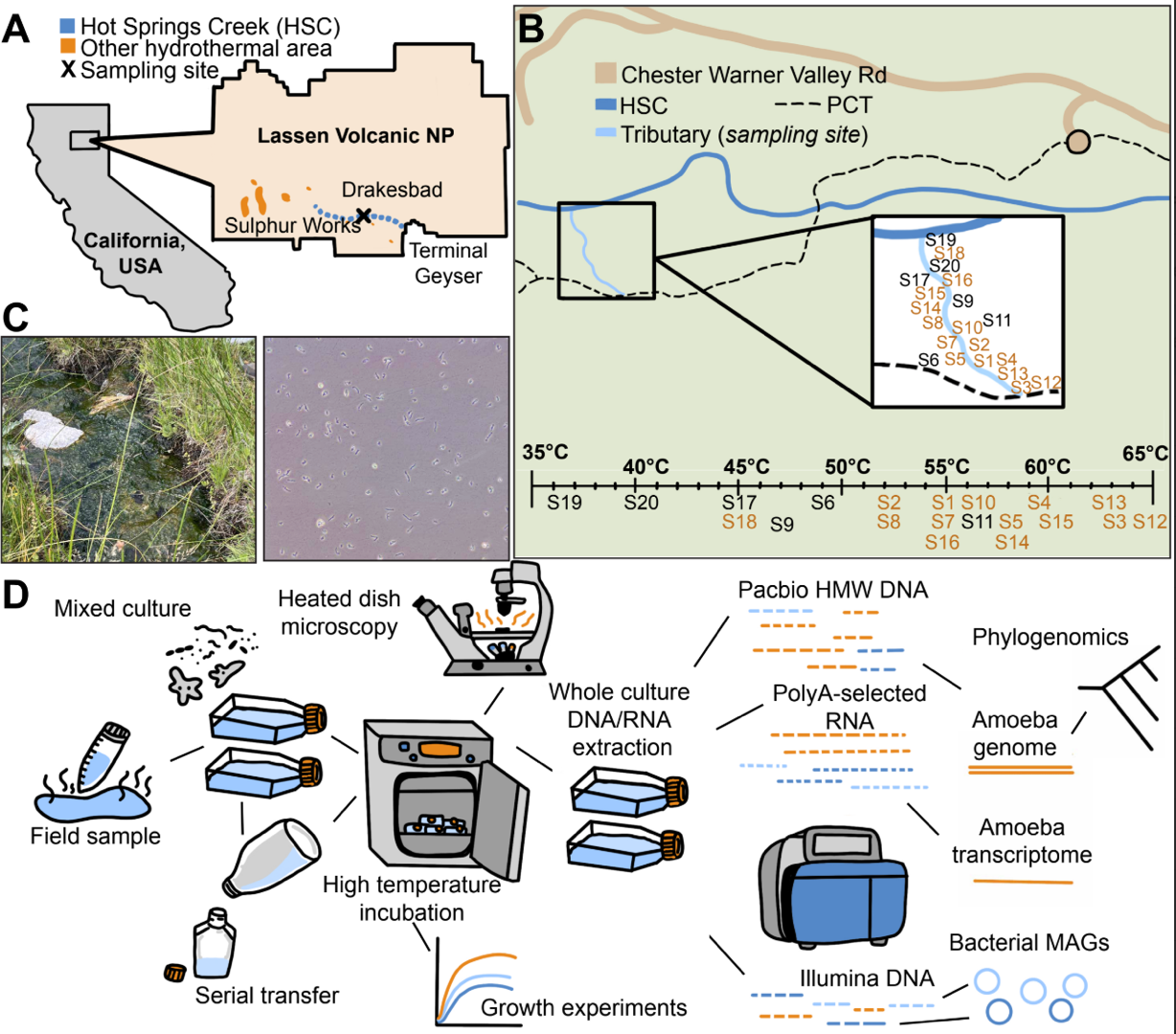

**Fig. S1. Overview of sampling site and study design.** **(A)** Sampling site is marked with an X off of Hot Springs Creek (HSC; dotted blue line) at Lassen Volcanic National Park. Orange denotes other hydrothermal areas near Hot Springs Creek. **(B)** Map of sampling site, depicting location of tributary (light blue) off of HSC (dark blue) along Chester Warner Valley Rd (brown) and the Pacific Crest Trail (dotted black). Zoom in of the tributary shows approximate locations of the 20 sampling sites above a line corresponding to temperature of each sampling site. Sites labeled in orange yielded cultures of *I. cascadensis* while sites labeled in black did not. **(C)** Left: Image of sampling site with prominent algal biofilm, surrounded by *Panicum* (panic grass). Right: Image of enrichment culture incubated at 57ºC observed at 10X magnification. **(D)** Overview of methods. We enriched amoebae from field samples via serial transfer resulting in a mixed culture of amoebae and bacteria. We incubated flasks at high temperature (57-60°C) to reach high amoeba density. We used established cultures for growth limit experiments and for heated dish microscopy. We extracted DNA from cultures for short and long read sequencing and RNA targeting amoeba transcriptomes.

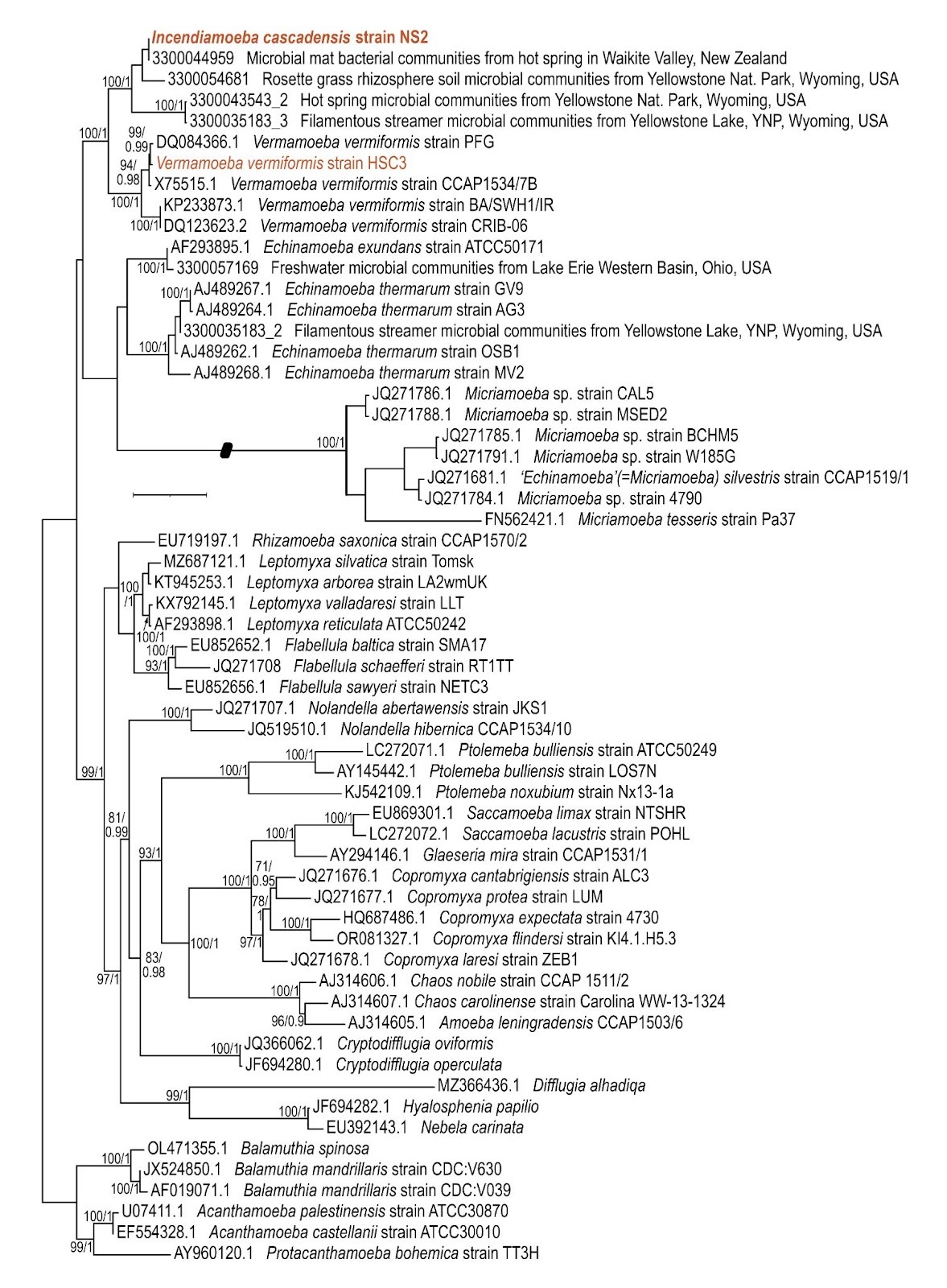

**Fig. S2. The 18S rRNA gene-based tree supports placement of *Incendiamoeba* as a novel genus and indicates global biogeographic distribution in geothermal habitats**.

18S rRNA tree built with IQ-TREE of *I. cascadensis* (bold, dark orange) and neighboring Tubulinea amoebae. Isolated *V. vermiformis* strain HSC3 highlighted in dark orange. Bootstrap support is provided.

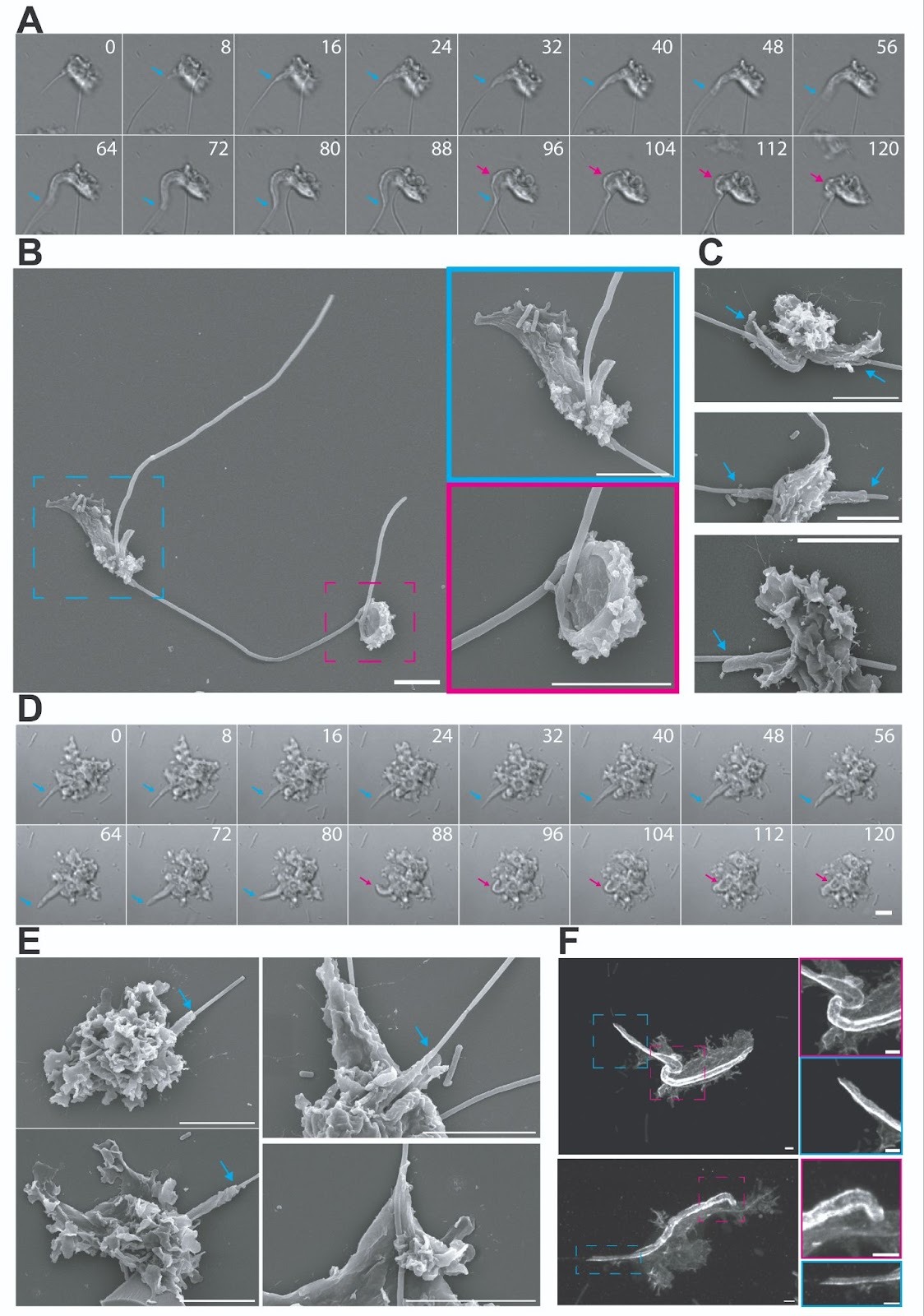

**Fig. S3. Feeding on filamentous bacteria appears to be facilitated by an initial tube-like structure which folds bacteria into the amoebae.** (**A)** Live cell imaging of *I. cascadensis* cells feeding at 60ºC. Tube-like structure is indicated by blue arrows, and the folded and enclosed bacterial cell (feeding cup) is indicated by a pink arrow. (**B-C**) SEM of feeding cells at 58ºC show similar structures to live cell imaging. Feeding tubes indicated by blue arrows/box, while folded cup bacterial structure is indicated by pink arrows/box, scale bars 5µm. (**D-F**) Feeding at 63ºC. (**D**) Live cell imaging (DIC) of *I. cascadensis* feeding at 63ºC. (**E**) SEM examples of feeding at 63ºC show feeding tubes (blue arrows). (**F**) Ultrastructure expansion microscopy confocal images of membrane labeling. Zoom-in of feeding tube (blue box) and folded bacterial ‘cup’ (pink box). Scale bars 5µm, not adjusted for expansion factor.

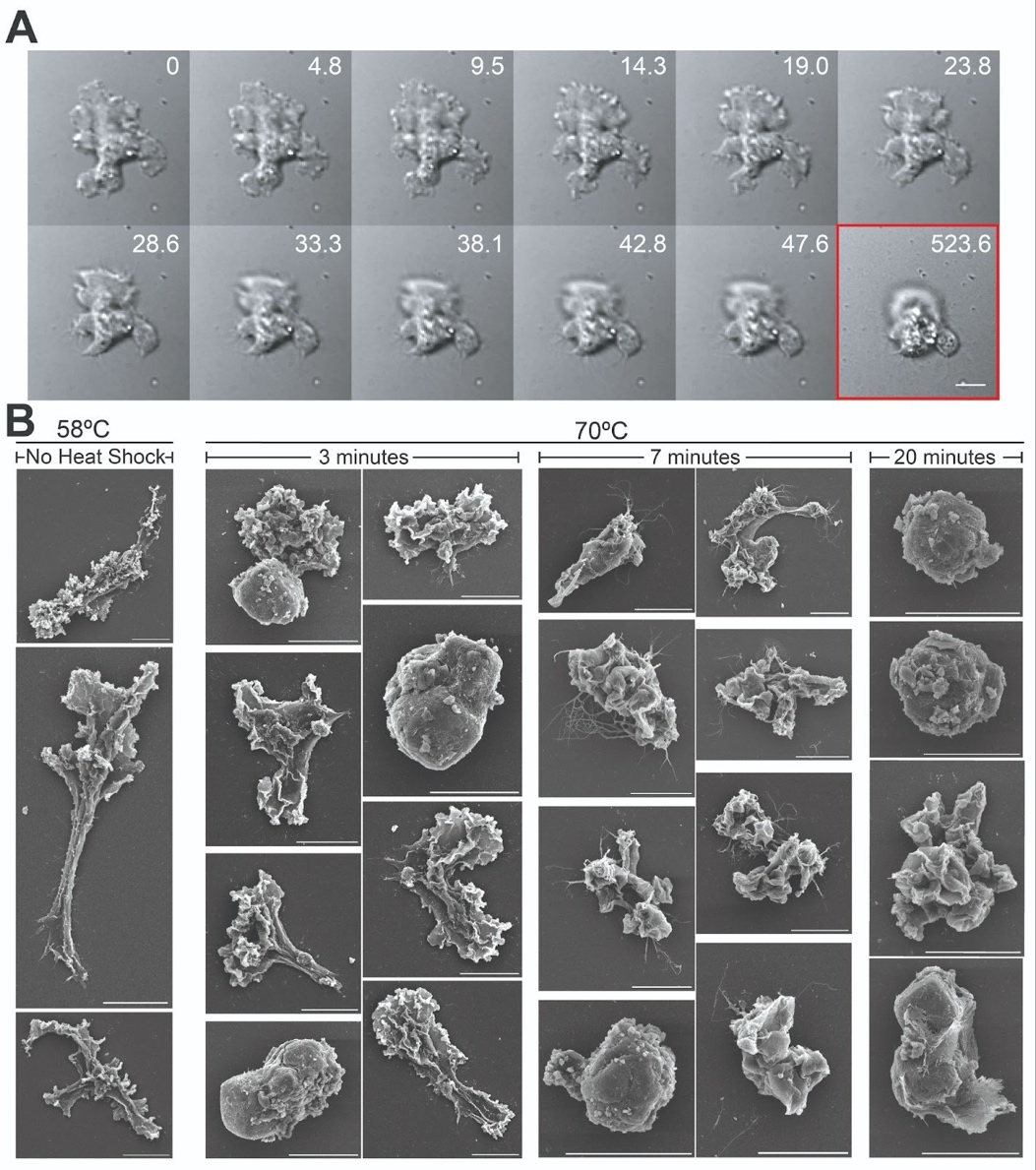

**Fig. S4. Rapid establishment of cysts in response to increasing temperatures as shown by live cell imaging and SEM.** (**A**) DIC timelapse of encystment at 70ºC. Time (seconds) is indicated in the upper right corner. The red box around the last frame indicates a jump in time increment. (**B**) SEM of cells at optimal temperature (left column) and heat shock at 70ºC for 3 minutes, 7 minutes, and 20 minutes.

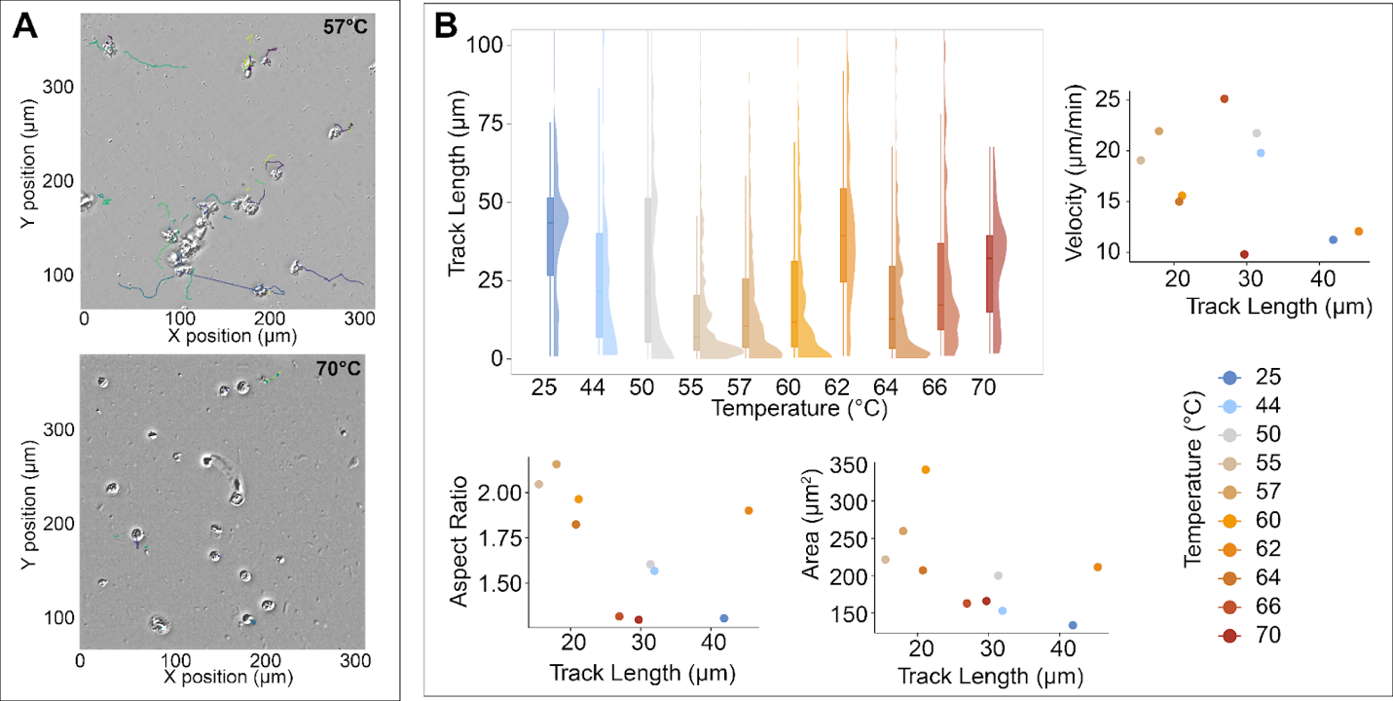

**Fig. S5. Quantifying tracking performance of time series imaging data.** (**A**) Example particle tracks from imaging at 57ºC (upper) and 70ºC (lower). Unique track identifier indicated by color. (**B**). Track length distribution across temperatures and comparison to extracted values such as velocity, aspect ratio, and area. Box and whisker plots show the interquartile range with median values as horizontal lines. Individual points are shown as a beeswarm distribution to illustrate spread of the data.

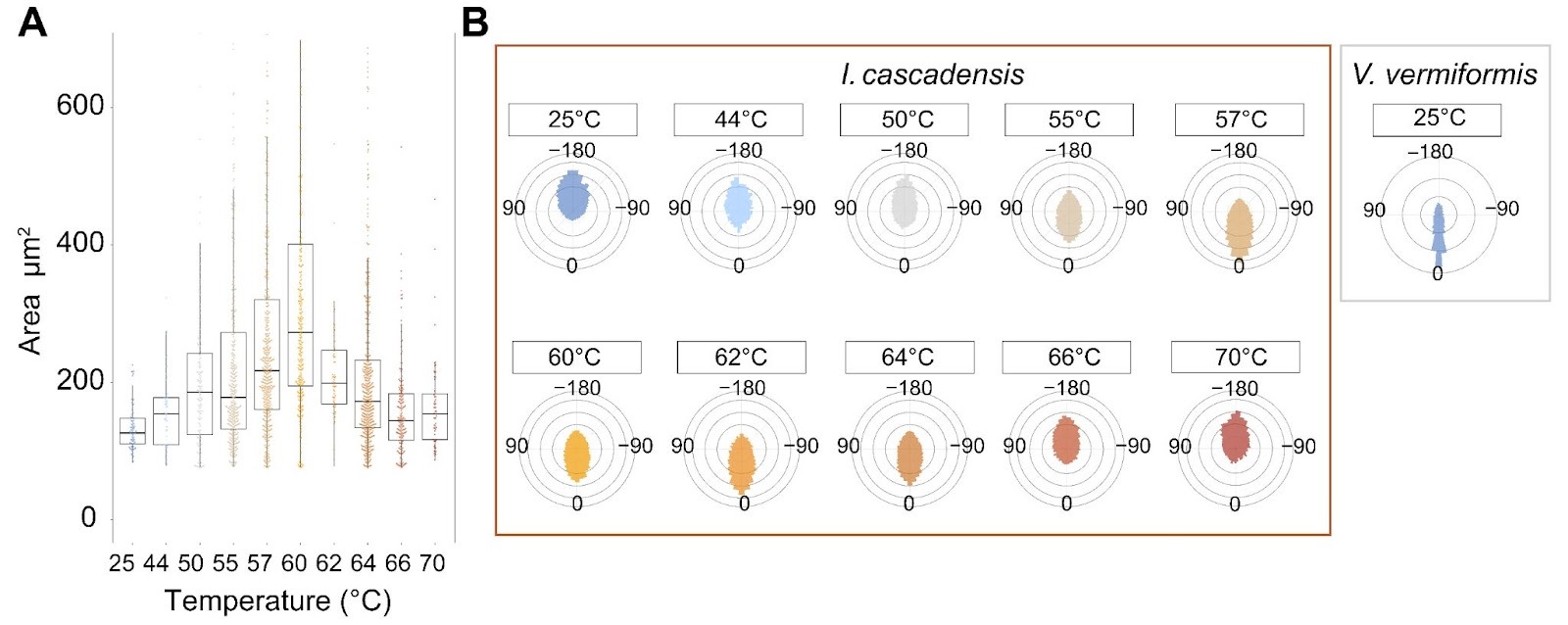

**Fig. S6. Cell area and directional changes provide additional support for active motility from 55ºC to 64ºC.** (**A**) Cell area across temperature. Box and whisker plots visualize the percentiles as follows: 75th (top edge), 25th (bottom edge), and median (bold line); whiskers extend to 1.5×IQR from the box, excluding outliers. Points represent individual cells per track. (**B**) Angle distributions plotted as polar histograms calculated from tracked cell centroids across temperature. Boxes around **B** plots are colored by species (*I. cascadensis* in dark orange and *V. vermiformis* in gray).

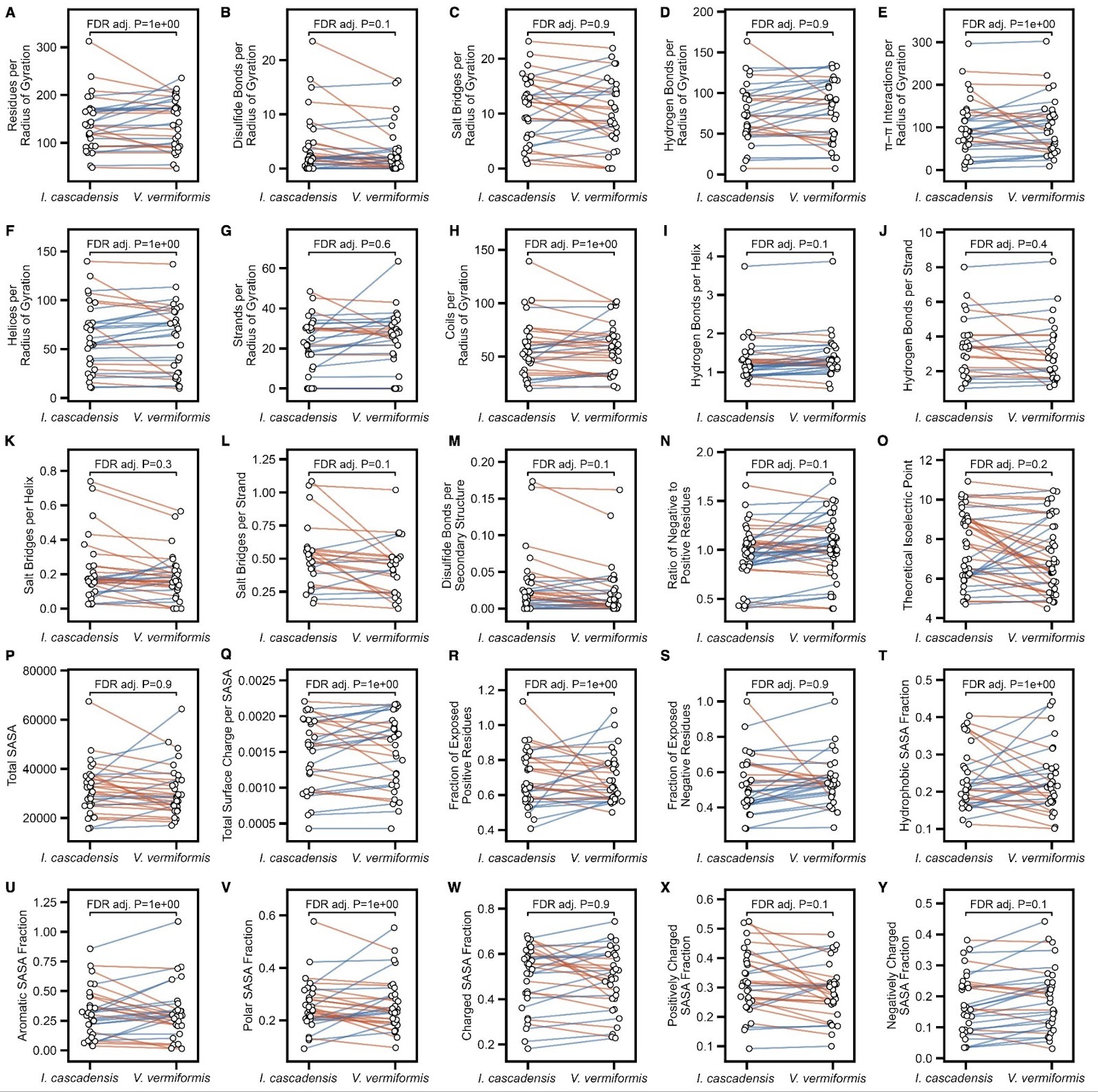

**Fig. S7. Predicted changes between analogous predicted proteins found in both *I. cascadensis* and *V. vermiformis* for a range of structurally derived metrics.** Metrics include (**A**) size, (**B-E**) intramolecular interactions, (**F-M**) secondary structure characteristics, (**N-O**) charge and (**P-Y**) solvent accessible surface area (SASA) for different groups of residues. Each dot represents a homologous protein found in both species, and each protein pair is connected by a line. Orange lines indicate a decrease from *I. cascadensis* to *V. vermiformis*, while blue lines indicate an increase. P-values are determined using a two-sided dependent T-test, with a BH correction for false discovery.

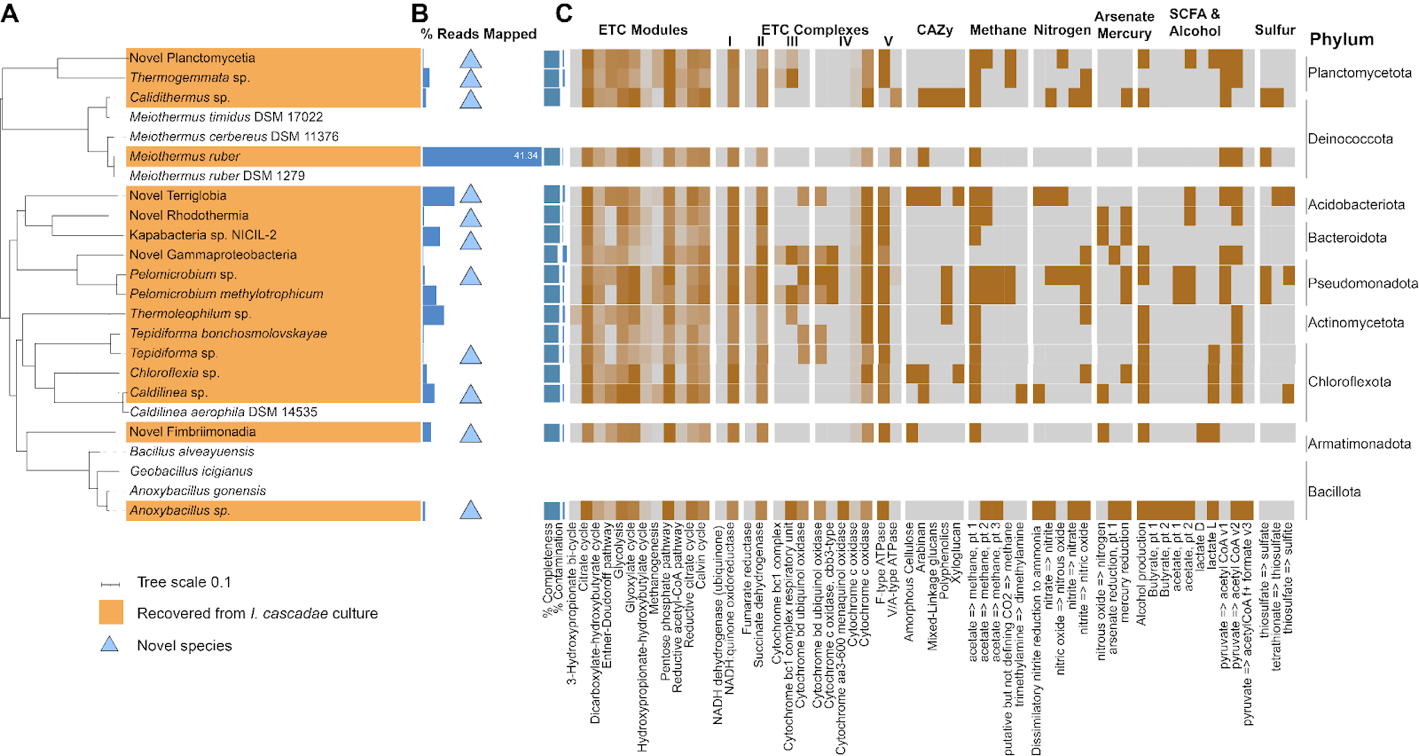

**Fig. S8. *Incendiamoeba cascadensis* grows with a phylogenetically and metabolically diverse consortium of bacteria.** (**A**) Phylogenetic tree of bacterial consortium and selected close relatives from a multiple sequence alignment. The most complete representative of each MAG recovered from multiple samples is included in the tree. Triangles indicate putative new species assigned by relative evolutionary divergence (RED) values. (**B**) Bars indicate percent reads of the concatenated set of reads from four samples mapped to each representative MAG. (**C**) Genome features of the representative MAGs including % completeness and contamination, as well as variable metabolic traits corresponding to modules and complexes of the electron transport chain (ETC), carbohydrate-active enzymes (CAZy), short chain fatty acid (SCFA) and alcohol production, methanogenesis and methanotrophy, and metabolism of nitrogen, arsenate, mercury, and sulfur.

**Supplementary Tables**

**Table S1. Metadata associated with all Hot Springs Creek tributary samples.**

Including sampling site code, year sampled, temperature, pH, whether *Incendiamoeba* was cultured from the site, and where metagenome or high molecular weight genome data were acquired.

**Table S2. Branch length comparisons and confidence interval assessment.**

Length to common ancestor (CA) for each species and average length to common ancestor (CA) for both species in within genus comparisons along with *Vermamoeba vermiformis* versus *Incendiamoeba* cascadensis. Length_to_CA represents raw branch lengths.

**Table S3. Morphometric data**. Major/Minor Axes (µm) averaged per particle track and area (µm²).

Table S4. Growth experiments. Population size (cells/mL) of individual flask replicates and average temperature readings of probes.

Table S5. Doubling summaries. Average estimated doubling time across batches, along with model statistics derived from log-transformed growth curves.

Table S6. Velocities. Median velocity (µm/min) and mean velocity binned by aspect ratio (µm/min). Velocity is reported as Interquartile Range (IQR), or 50% of the values, to highlight typical speeds and de-emphasize outliers.

**Table S7. Motility stats.** Mean square displacement (MSD) slope **(**α) and displacement ratio, convex hull area**.**

**Table S8. Cell Shape Switching Rates.** Cell shape binned by aspect ratio where values greater than 1.2 but less than 2.5 are categorized as "amoebiform" while 2.5 or greater are categorized as "vermiform". Switching rates are reported as the average number of switches per minute and their reciprocal, average time for each switch. Standard error of the mean (SEM) is used to report variability.

**Table S9**. **Genome assembly statistics across Amoebozoa genomes included in gene content analysis.** Total genome length, percent GC, BUSCO completeness and contamination, number of contigs, and N50 score.

**Table S10. KEGG functional comparison.** Enrichment comparison of KEGG functions between two *I. cascadensis* genomes and three mesophilic Amoebozoa. Average excess is the number of species with more or fewer copies of an ortholog than the two *I. cascadensis*.

**Table S11. Comparative physicochemical properties between *I. cascadensis* and *V. vermiformis* homologous proteins.** Size, intramolecular interactions, secondary structure characteristics, charge, and solvent accessible surface area of 30 homologous proteins between *I. cascadensis* and *V. vermiformis*.

**Table S12. Bacterial MAG stats and metabolic predictions.** Taxonomy of bacterial MAGs recovered from *Incendiamoeba* cultures, percent of reads mapped, completeness, contamination, and DRAM functions.

**Other Supplementary Materials**

**Movie S1. Attachment of *I. cascadensis* in amoebiform state.**

DIC time series on left, RICM on right. Elapsed time 6 minutes 24 seconds, scale bar 5µm.

**Movie S2. Attachment of *I. cascadensis* in vermiform state.**

DIC time series on left, RICM on right. Elapsed time 55 seconds, scale bar 5µm.

**Movie S3. Attachment of *Vermamoeba vermiformis* in amoebiform and vermiform states.**

DIC time series on left, RICM on right. Elapsed time 1 minute 26 seconds, scale bar 5µm.

**Movie S4. Novel amoeba motility suggests nucleus-first migration.**

Nuclear (NucBlue; colored blue) and actin (SiR-actin; colored red) stained cells at 60ºC. Elapsed time 11 minutes 41 seconds, scale bar 20µm.

**Movie S5. Feeding on filamentous bacteria.**

*I. cascadensis* feed on filamentous bacterial biofilms. Elapsed time 25 minutes, scale bar 10µm.

**Movie S6. Feeding on filamentous bacteria close-up**.

Movie of *I. cascadensis* cell feeding from Fig. S3D, elapsed time 8 minutes 48 seconds, scale bar 5µm.

**Movie S7. 44ºC motility.**

Example acquisition of cells incubated at 44ºC. DIC time series. Elapsed time 5 minutes, scale bar 20µm.

**Movie S8. 57ºC motility.**

Example acquisition of cells incubated at 57ºC. DIC time series. Elapsed time 5 minutes, scale bar 20µm.

**Movie S9. 64ºC motility.**

Example acquisition of cells incubated at 64ºC. DIC time series. Elapsed time 5 minutes, scale bar 20µm.

**Movie S10**. **Encystment from heat shock.**

Starting at 60ºC, cells display normal motility. At 3 minutes 39 seconds, cells are exposed to 70ºC media for 10 seconds before a return to 60ºC (temperature labeled on screen). Total time 7 minutes, 1 second intervals, scale bar 20µm.

**Movie S11. Encystment close-up showing hairlike structures.**

Single *I. cascadensis* cell encysting with hairlike structures emerging. Elapsed time 3 minutes 57 seconds, scale bar 5µm.

**Movie S12. Novel amoeba displays two motility modes at high temperatures.**

DIC time series at 100X with Bioptechs heated chamber reveals vermiform and amoebiform movement alternate in cells at 60ºC. Elapsed time: 9 minutes 55 seconds.
